# Swimming motility and chemotaxis control the spatial organization, persistence, and inflammatory activity of a model intestinal pathobiont

**DOI:** 10.1101/779983

**Authors:** Travis J. Wiles, Brandon H. Schlomann, Elena S. Wall, Reina Betancourt, Raghuveer Parthasarathy, Karen Guillemin

**Author notes:** These authors contributed equally (Travis J. Wiles, Brandon H. Schlomann).

## Abstract

Understanding the processes that spatially restrict resident gut bacteria and the mechanisms by which disease-causing pathobionts escape this control will open new avenues for microbiome-based therapies. Using live imaging and genetically engineered bacteria, we discovered that flagella-based swimming motility and chemotaxis enable a model *Vibrio* pathobiont to govern its own spatial organization within the larval zebrafish gut and to persist in the face of the disruptive forces of intestinal flow. Bacterial mutants lacking motility traits became aggregated and lumenally confined, making them susceptible to periodic expulsion from the host. Consequently, non-motile and non-chemotactic mutants experienced large fluctuations in absolute abundance and impaired interbacterial competition. Further, we found that motile bacterial cells induce expression of the proinflammatory cytokine TNFα in gut-associated macrophages and the liver. Using inducible genetic switches, we demonstrate that swimming motility can be manipulated in situ to modulate the spatial organization, persistence, and inflammatory activity of gut bacteria.

## INTRODUCTION

Bacterial pathobionts are indigenous members of the microbiome that have a latent ability to undermine host health. Although pathobionts have been implicated in numerous diseases (Chow et al., 2011; Hajishengallis and Lamont, 2016), the factors underlying their pathogenic potential remain poorly defined. Understanding how the host constrains the virulent activities of resident bacteria and the mechanisms pathobionts use to escape this control will lead to new microbiome-based therapies for improving human and animal health.

One way the host keeps pathobionts in check within the intestine is by imposing restrictions on microbiota spatial organization. The most recognized spatial control measures employed by the host are mucus, immunoglobulins, and antimicrobial peptides, which confine bacteria to the intestinal lumen, away from mucosal surfaces (Cullender et al., 2013; Johansson et al., 2013; Vaishnava et al., 2011). In turn, it is thought that intense competition for resources pushes bacteria to evolve strategies that allow them to subvert host control and exploit opportunities to occupy new spatial niches (Finlay and Falkow, 1989; Foster et al., 2017). In line with this idea, several prototypic pathobionts undergo blooms in abundance that are coincident with shifts in intestinal biogeography (Carvalho et al., 2012; Chow et al., 2011; Gevers et al., 2014; Kostic et al., 2014). A potential trait underlying this behavior that is common to many pathobionts—as well as numerous bona fide pathogens—is flagella-based swimming motility (Chaban et al., 2015; Elhenawy et al., 2019; Ottemann and Miller, 1997). Swimming motility gives bacteria the agency to govern their own spatial organization and access niches that enhance growth and survival (Raina et al., 2019; Wei et al., 2011; Yawata et al., 2014). In some cases, it has been shown that flagellin, the protein subunit comprising the bacterial flagellum, is a major driver of pathobiont-induced inflammation (Ayres et al., 2012). Further underscoring the relationship between pathobionts and motility, mouse studies have revealed that hosts have several mechanisms for detecting and quenching flagellar motility that are critical to intestinal homeostasis (Ayres et al., 2012; Cullender et al., 2013; Fulde et al., 2018; Okumura et al., 2016). However, despite progress in understanding the pathogenic consequences of bacterial motility for the host, much remains unknown about how motility behaviors promote intestinal colonization and provide pathobionts with a competitive advantage.

Answers to these questions have started to emerge from our studies of how diverse bacterial taxa colonize the zebrafish intestine. The optical transparency and small size of larval zebrafish make them an ideal vertebrate model for probing how bacteria use motility to spatially organize their populations within a living animal. With light sheet fluorescence microscopy (LSFM) it is possible to capture the full three-dimensional architecture of bacterial populations at single bacterial cell resolution across the entire length of the larval intestine (Parthasarathy, 2018). In addition, the spatiotemporal dynamics of bacterial and host cells can be followed in real time or over the course of many hours. Using LSFM, we have found that for many non-inflammatory commensal bacteria native to the zebrafish microbiome, the bulk of their populations are non-motile and reside as dense multicellular aggregates within the intestinal lumen (Schlomann et al., 2018; Wiles et al., 2018). Notably, this pattern of bacterial spatial organization is consistent with histological data from both the mouse and human intestine (Swidsinski et al., 2005, 2007; van der Waaij et al., 1996; Welch et al., 2017). We discovered that in this aggregated regime, bacteria are extremely vulnerable to intestinal flow. Consequently, aggregated bacterial populations can be stochastically expelled from the host in large numbers, producing punctuated drops in abundance (Schlomann et al., 2019; Wiles et al., 2016).

In contrast, unlike most zebrafish gut bacteria studied thus far, we have identified an isolate of non-toxigenic *Vibrio cholerae* (strain ZWU0020, further referred to as “*Vibrio*” for brevity) that exhibits pathobiont-like characteristics and assembles intestinal populations made up of planktonic cells displaying vigorous swimming motility (Rolig et al., 2017; Wiles et al., 2016). The mass swimming behavior of *Vibrio* populations gives them a liquid-like spacing-filling property that promotes frequent and close contact with the intestinal mucosa (Wiles et al., 2016). This attribute appears to make *Vibrio* highly resistant to intestinal expulsion. As a result, *Vibrio* stably colonizes the intestine and reaches absolute abundances that are up to ten times higher than other zebrafish symbionts (Schlomann et al., 2019). *Vibrio*’s unique intestinal lifestyle is also potentially linked to its pathobiont character, which is marked by its ability to supplant established, naturally aggregated bacterial populations (Wiles et al., 2016), and induce intestinal inflammation and exacerbate pathology in susceptible hosts (Rolig et al., 2015, 2017).

In the present work we set out to identify the mechanisms by which *Vibrio*’s motility behaviors control its ability to colonize the intestine and contribute to its proinflammatory potential. We found that *Vibrio* specifically requires sustained swimming motility to resist intestinal flow and persist at high abundances. Mutants lacking either the flagellar motor or chemotaxis are attenuated for colonization and interbacterial competition. Using mutant animals with decreased intestinal transport and bacteria carrying inducible genetic switches that allow motility behaviors to be toggled in situ, we demonstrate that loss of motility or chemotaxis leads to increased aggregation and expulsion. Further, colonizing transgenic animals encoding a fluorescent reporter of tumor necrosis factor alpha (TNFα) expression, which is a proinflammatory cytokine, revealed that *Vibrio* is capable of inducing inflammation both locally within the intestine and systemically, particularly in cells of the liver. We found that *Vibrio*’s proinflammatory activity is dependent on motility and chemotaxis, and that macrophages associated with the intestine are a major cell type that responds to bacterial motility. Finally, activating motility in established, non-motile populations showed that host tissues are remarkably sensitive to sudden changes in bacterial motility and spatial organization.

Our work yields mechanistic insights into the form and function of the intestinal ecosystem. We have disentangled the requirements for motility behaviors of a model pathobiont during multiple stages of host colonization. Moreover, we identified that a motility-based intestinal lifestyle has potent proinflammatory consequences, emphasizing that bacterial swimming motility and the host countermeasures that control it are potential targets for therapeutic manipulation of the microbiome.

## RESULTS

### Loss of swimming motility or chemotaxis attenuates intestinal colonization and interbacterial competition

To dissect the role of flagellar motility during intestinal colonization, we generated two motility-deficient *Vibrio* mutants (Figure S1A). To test swimming motility in general, we deleted the two-gene operon *pomAB* that encodes the polar flagellar motor (creating Δmot). To test *Vibrio*’s ability to spatially organize its populations in response to environmental cues, we deleted the gene *cheA2*, which encodes a histidine kinase necessary for chemotaxis (creating Δche). In vitro, Δmot exhibited complete loss of swimming motility whereas Δche had a run-biased behavior with swim speeds comparable to wild type but failed to chemotax in soft agar (Figure S1B). Both motility mutants displayed normal growth and assembled a single polar flagellum similar to wild type (Figure S1C and S1D).

We first assessed the absolute abundances of each strain over time by gut dissection and cultivation. We inoculated wild-type *Vibrio*, Δmot, and Δche individually into the aqueous environment of four-day-old germ-free larval zebrafish (Melancon et al., 2017). *Vibrio* rapidly colonized germ-free animals to high abundance, reaching a maximal carrying capacity of 10^5^– 10^6^ cells per intestine by 24 h post-inoculation (hpi) and maintaining a high-level of abundance through 72 hpi (Figure 1A). In contrast, Δmot and Δche displayed attenuated intestinal colonization phenotypes (Figure 1A). Both mutants were slow to access the zebrafish intestine, suggesting that they have reduced immigration rates, and reached maximal abundances at 24 hpi that were 10–100-fold lower than wild type (Figure 1A).

**Figure 1.**
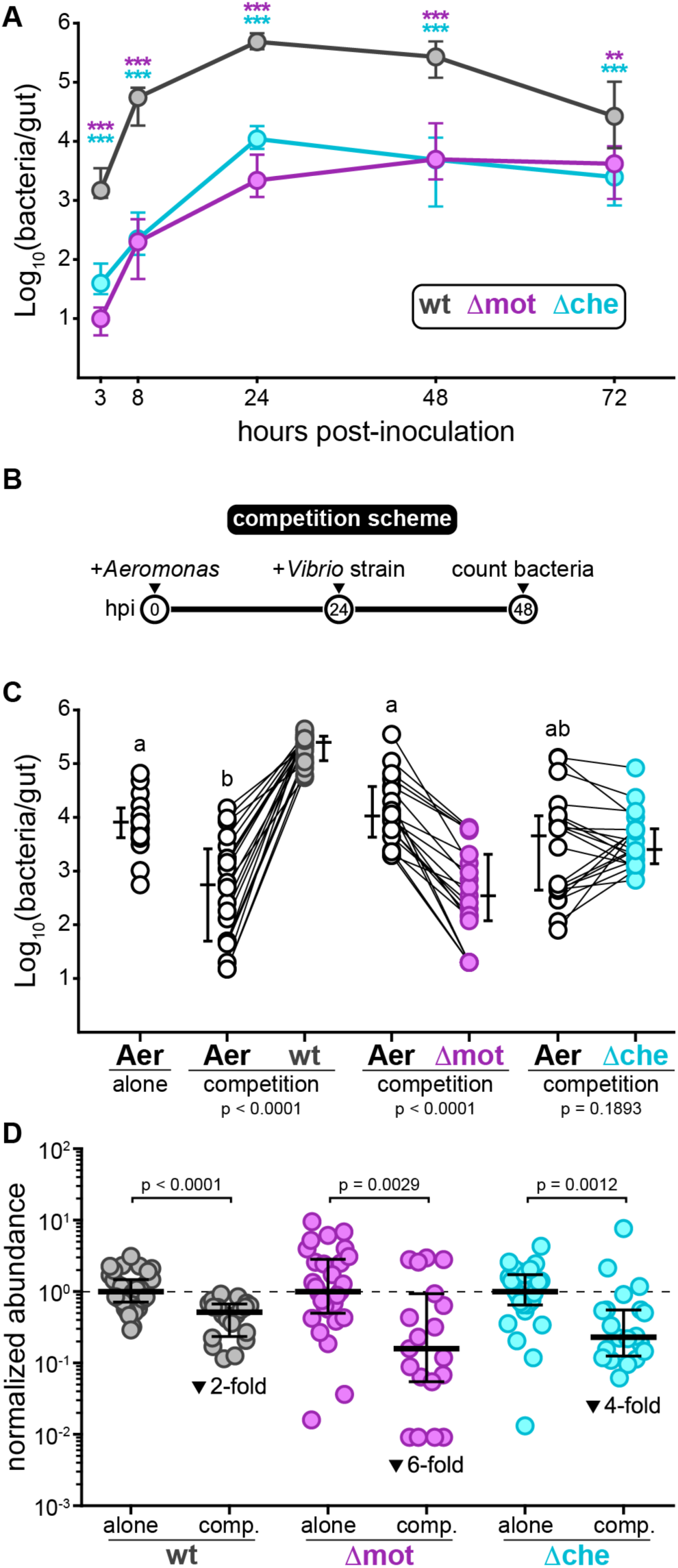
Loss of swimming motility or chemotaxis attenuates intestinal colonization and interbacterial competition. (see also Figure S1) **(A)** Abundances of wild-type *Vibrio*, Δmot, and Δche during mono-association. Plotted are medians and interquartile ranges (n ≥ 17 animals/marker). Significant differences between each mutant and wild-type determined by Mann-Whitney (purple asterisks: Δmot; cyan522 asterisks: Δche). ***p < 0.0001, **p = 0.0002. **(B)** Experimental timeline of *Aeromonas*–*Vibrio* competition. **(C)** Abundances of *Aeromonas* while alone or in competition with wild-type or mutant *Vibrio*s. Letters denote significant differences between *Aeromonas* treatments. p < 0.05, Kruskal-Wallis and Dunn’s. Adjacent bars denote medians and interquartile ranges. Significant differences based on Wilcoxon between *Aeromonas* and each *Vibrio* strain are noted below each competition. **(D)** Abundances of wild-type and mutant *Vibrio*s during competition with *Aeromonas* (from panel C) normalized to abundances during mono-association at 24 hpi (from panel A). Bars denote medians and interquartile ranges. Significant differences determined by Mann-Whitney. Fold-decreases based on medians.

We next compared the ability of wild-type *Vibrio* and each mutant to invade an established population of *Aeromonas veronii* (strain ZOR0001, further referred to as “*Aeromonas*”). Like *Vibrio*, *Aeromonas* species are abundant members of the zebrafish intestinal microbiota (Stephens et al., 2016). Previous studies suggest that these two genera naturally compete against one another within complex intestinal communities (Phelps et al., 2017). In addition, we have shown that *Vibrio* is capable of invading and displacing established *Aeromonas* populations in gnotobiotic animals (Wiles et al., 2016). Following the competition scheme depicted in Figure 1B, we found that each *Vibrio* strain had a distinct competitive interaction with *Aeromonas* (Figure 1C). Wild-type *Vibrio* potently colonized *Aeromonas*-occupied intestines and induced 10–100-fold drops in *Aeromonas* abundances (Figure 1C). Zebrafish colonized with the Δmot mutant, however, were dominated by *Aeromonas*, which did not experience any significant declines in abundance compared to mono-association (Figure 1C). Invasion with the Δche mutant had an intermediate impact on *Aeromonas* abundances and the two appeared to co-exist (Figure 1C). Comparing abundances during competition to those during mono-association showed that each *Vibrio* strain’s colonization was hindered to varying degrees while invading established *Aeromonas* populations (Figure 1D). Wild-type *Vibrio* abundances were only 2-fold lower during competition than during mono-association (Figure 1D). In contrast, Δmot abundances were 6-fold lower (and in several instances reduced by up to 100-fold), whereas the impact on Δche abundances was intermediate with a 4-fold reduction (Figure 1D). Overall, these data show that *Vibrio* requires swimming motility and chemotaxis for normal intestinal colonization and interbacterial competition.

### Motility and chemotaxis mutants have altered intestinal spatial organization

Wild-type *Vibrio* cells strongly localize to the larval zebrafish foregut (Figure 2A and 2B) (Schlomann et al., 2018), which is an anatomical region comparable to the mammalian small intestine (namely, the duodenum and jejunum) (Lickwar et al., 2017; Wang et al., 2010), and display a highly active swimming behavior both within the intestinal lumen and at mucosal surfaces (Wiles et al., 2016). To determine how motility and chemotaxis contribute to *Vibrio*’s cellular behavior and spatial organization within the intestine, we examined wild type, Δmot, and Δche in live animals using LSFM. A fluorescently marked variant of each strain was first mono-associated with germ-free animals and then imaged at 48 hpi. As expected, wild-type *Vibrio* assembled dense populations concentrated within the foregut that were almost entirely composed of planktonic cells swimming in the lumen as well as within the intestinal folds (Figure 2C–2G, Movies S1 and S2). In contrast, Δmot and Δche assembled populations with greatly altered behavior and spatial organization.

**Figure 2.**
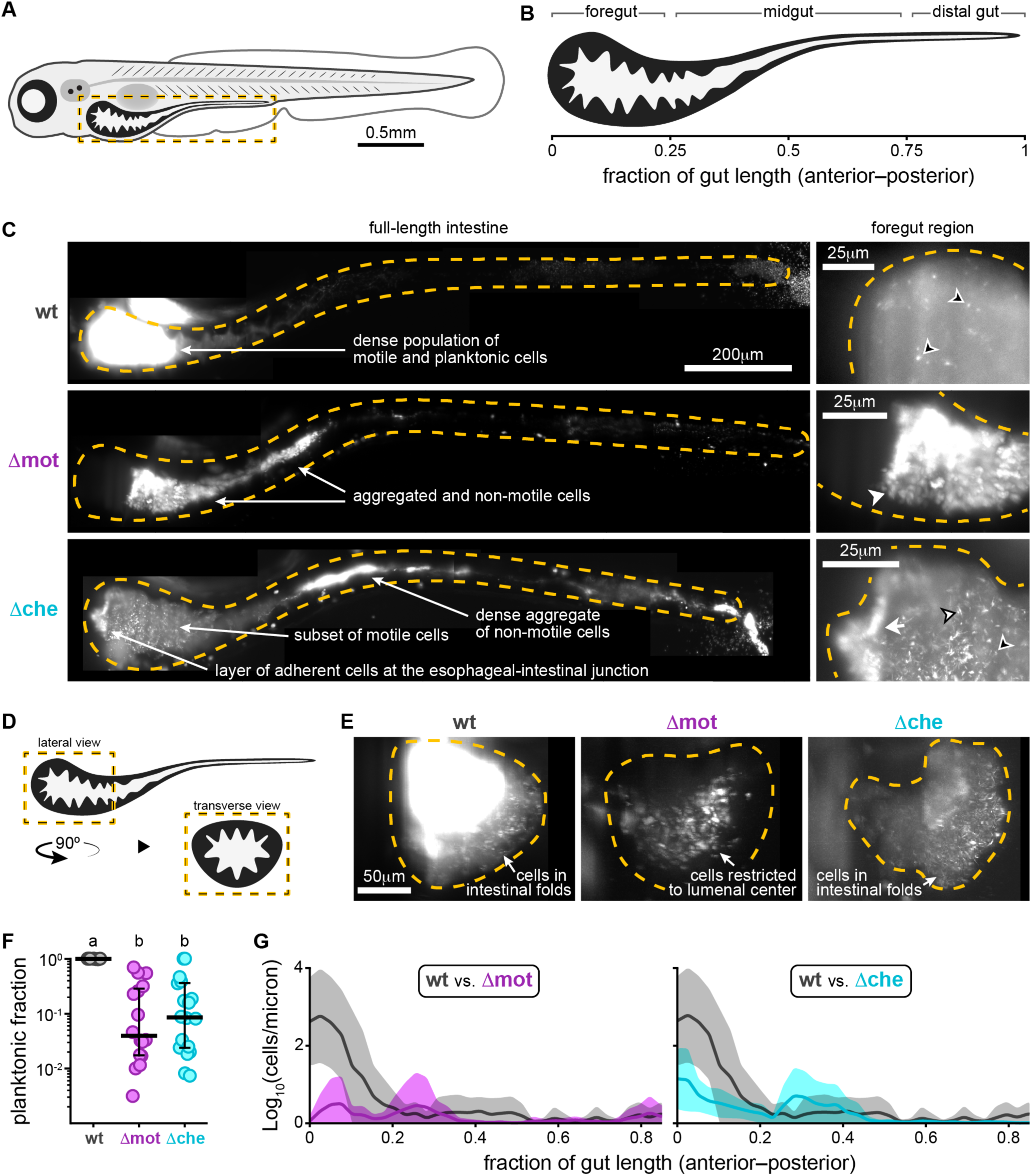
Motility and chemotaxis mutants have altered intestinal spatial organization. (see also Figure S1) **(A)** Cartoon of a 6-day-old zebrafish. Dashed box marks intestinal region imaged by LSFM. **(B)** Anatomical regions of the larval zebrafish intestine. **(C)** Maximum intensity projections acquired by LSFM showing the spatial organization of wild-type *Vibrio* (top), Δmot (middle), and Δche (bottom) within the intestine. Top right inset shows a zoomed-in view of wild-type *Vibrio* cells in a separate fish that was colonized with a 1:100 mixture of green- and red-tagged variants so that the cellular organization of the dense *Vibrio* population could be discerned. The dilute channel (green) is shown. Dashed lines mark approximate intestinal boundaries. Open arrowheads: single bacterial cells; solid arrowheads: small aggregates; tailed arrowheads: large aggregates. Arrowheads with a black stroke mark swimming cells, which appear as comet-like streaks. **(D)** Cartoon showing the intestinal region pictured in panel (E) **(E)** Maximum intensity projections acquired by LSFM showing transverse view of the foregut region colonized with wild type, Δmot, or Δche. **(F)** Fraction of planktonic cells contained within each strain’s population. Each circle is a measurement from a single intestinal population. Bars denote medians and interquartile ranges. Letters denote significant differences. p < 0.05, Kruskal-Wallis and Dunn’s. **(G)** Image-derived abundances of wild type (n = 7), Δmot (n = 4), and Δche (n = 5) with respect to position along the length of the gut. Shaded regions mark confidence intervals.

Populations of Δmot were non-motile whereas Δche had a small subset of motile cells that could often be observed in the foregut (Figure 2C, Movie S1). Unexpectedly, both Δmot and Δche became highly aggregated within the intestine (Figure 2C–2E) despite exhibiting no signs of aggregation in vitro (Figure S1). The fraction of planktonic cells contained within each mutant population was >10-fold lower than wild type (Figure 2F). The aggregated cells of Δmot appeared to be mostly restricted to the lumen whereas the swimming cells of Δche, like wild type, were observed within the intestinal folds (Figure 2D and 2E, Movie S2). The Δmot mutant was also largely excluded from the anterior most portion of the foregut whereas Δche often formed an adherent layer of cells on the anterior wall near the esophageal-intestinal junction (Figure 2C). The population-wide aggregation of both mutants (which we refer to as cohesion) coincided with an overall posterior shift in distribution compared to wild type (Figure 2C and 2G). This shift in distribution is consistent with previous findings of strong correlations across bacterial species between cohesion and localization along the zebrafish intestine (Schlomann et al., 2018). In total, our live imaging data show that *Vibrio* requires swimming motility and chemotaxis to spatially organize its populations within the intestine. Further, Δmot and Δche formed aggregated and lumen-restricted populations reminiscent of other zebrafish bacterial symbionts, like *Aeromonas*, that largely lack swimming motility in vivo (Schlomann et al., 2018; Wiles et al., 2018).

### Swimming motility and chemotaxis promote persistence by enabling bacteria to counter intestinal flow and resist expulsion

We previously found that naturally aggregated bacteria are vulnerable to intestinal flow and expulsion from the host (Schlomann et al., 2019; Wiles et al., 2016). To explore if the attenuated colonization phenotypes of Δmot and Δche are connected to their perturbed spatial organization in a way that causes increased sensitivity to the intestine’s mechanical forces, we followed the spatiotemporal dynamics of wild-type *Vibrio* and each mutant in live animals by LSFM. Prior to imaging, each strain was given 24 h to reach its respective carrying capacity in germ-free zebrafish. Despite wild-type *Vibrio* showing modest declines in abundance from 24–72 hpi (Figure 1A), it was highly uniform and stable over periods of >10 h, maintaining its abundance, low cohesion, and foregut localization (Figure 3A, Movie S3). We note that image-based quantification of wild-type *Vibrio* abundances was performed in a concurrent study (Schlomann et al., 2019) and have been replotted here. In contrast, Δmot and Δche underwent dramatic fluctuations in their abundances and spatial organization (Figure 3A, Movie S3). Cells and small aggregates in Δmot and Δche populations appeared to become packed by intestinal contractions into large masses within the midgut before being abruptly expelled. Autofluorescent material was often observed surrounding aggregated cells, suggesting that host mucus was involved in this process. Image-based quantification of absolute abundances showed that >90% of Δmot and Δche populations could be lost in a single collapse event (Figure 3A). Following collapses, residual small aggregates in the midgut and low numbers of planktonic cells in the foregut appeared to undergo bursts in replication that effectively restored the abundance and spatial organization of the population before the next collapse. Animating the relationship between cohesion and intestinal localization for each *Vibrio* strain across animals over time showed that both mutant populations exhibit large fluctuations in spatial organization whereas wt *Vibrio* populations are highly stable (Movie S4).

**Figure 3.**
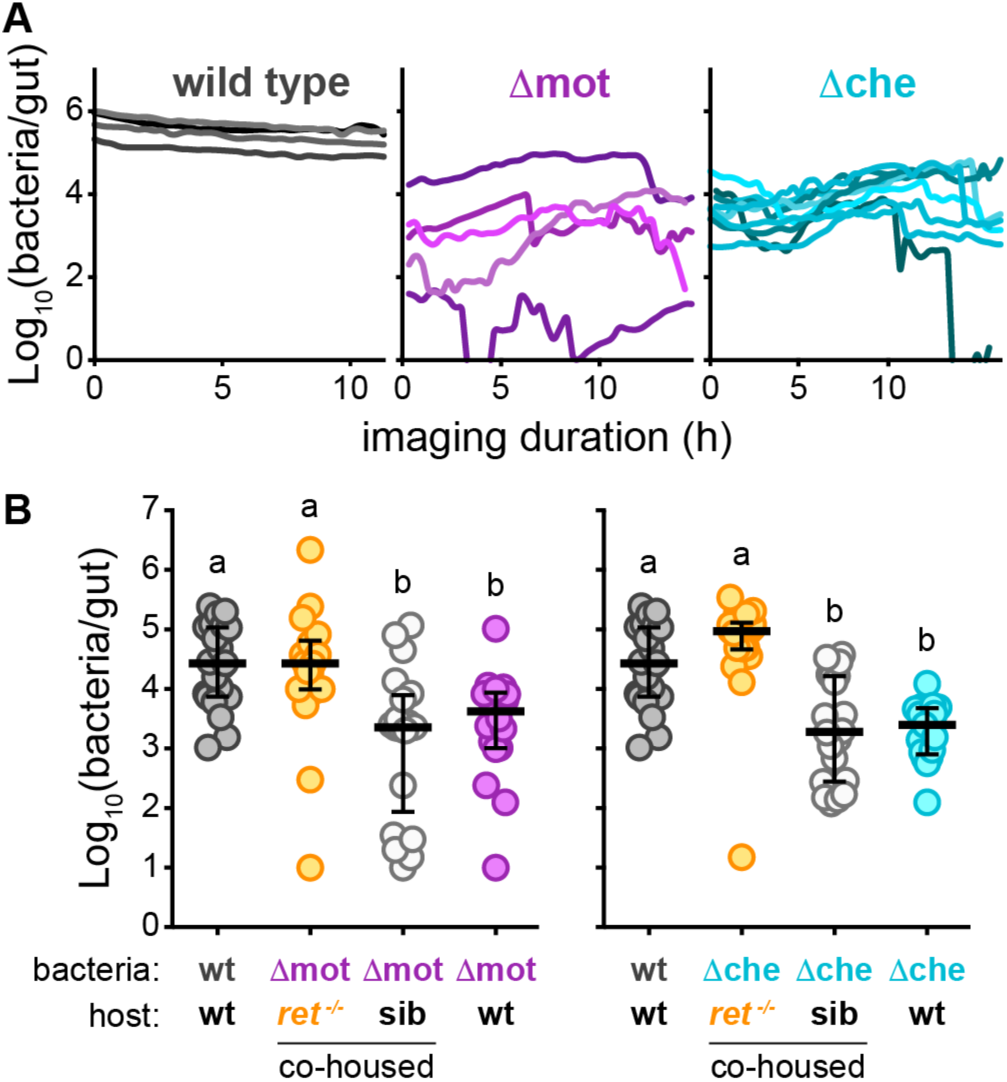
Swimming motility and chemotaxis promote persistence by enabling bacteria to counter intestinal flow and resist expulsion. (see also Figure S2) **(A)** Image-based quantification of abundances over time for wild-type *Vibrio*, Δmot, and Δche. Lines represent individual populations in individual fish. **(B)** Cultivation-based quantification of abundances for Δmot and Δche in co-housed *ret*^-/-^ mutant hosts and wild-type/heterozygous sibling controls (sib). Abundances of wild-type *Vibrio*, Δmot, and Δche in wild-type hosts (from Figure 1A, 72 hpi) are shown for comparison. Bars denote medians and interquartile ranges. Letters denote significant differences. p < 0.05, Kruskal-Wallis and Dunn’s.

Our live imaging results suggested that the altered spatial organization of Δmot and Δche populations, namely their increased cohesion, makes them more susceptible to intestinal flow and expulsion, and thus is likely the cause of their reduced abundances. This putative mechanism contrasts with the general assumption that swimming motility and chemotaxis primarily promote bacterial growth by facilitating nutrient foraging and avoidance of hostile environments. To probe the likelihood of these two different mechanisms we quantified the in vivo growth rates of Δmot and Δche (see Methods). We found that both Δmot and Δche exhibit intestinal growth rates (Δmot = 0.7 ± 0.3 hr^-1^ [n = 2]; Δche = 0.9 ± 0.4 hr^-1^ [n = 5]) that are comparable to a previously determined wild-type *Vibrio* growth rate of 0.8 ± 0.3 hr^-1^ (mean ± standard deviation) (Wiles et al., 2016). This result supports the idea that the reduced intestinal abundances of Δmot and Δche are not due to attenuated growth, but rather are a consequence of altered behavior and spatial organization that increases susceptibility to intestinal flow and expulsion.

To test this expulsion-based mechanism more directly, we assessed whether the abundance of Δmot and Δche could be rescued in *ret*^-/-^ mutant zebrafish hosts, which have reduced intestinal transport due to a dysfunctional enteric nervous system (Ganz et al., 2018; Wiles et al., 2016). Humans with *ret* mutations can develop Hirschsprung Disease, which is an affliction characterized by intestinal dysmotility and altered gut microbiome composition (Gosain and Brinkman, 2015; Heanue and Pachnis, 2007). Strikingly, we found that the intestinal abundances of both Δmot and Δche were fully rescued to wild-type levels in *ret*^-/-^ mutant animals (Figure 3B). In contrast, Δmot and Δche abundances in co-housed sibling control animals mirrored those in wild-type animals (Figure 3B). Importantly, we found that wild-type *Vibrio* shows no change in intestinal abundance in *ret*^-/-^ mutant animals (Figure S2A), indicating that the rescue of Δmot and Δche is not due to a general overgrowth phenomenon. In addition, inspecting the spatial organization of Δmot in *ret*^-/-^ mutant animals revealed that in some instances Δmot populations displayed relocalization to the anterior portion of the foregut, suggesting that intestinal flow is responsible for Δmot’s posterior-shifted distribution in wild-type animals (Figure S2B). Together, these results provide further evidence that swimming motility and chemotaxis can act as a mechanism for resisting intestinal flow to promote intestinal persistence.

### Sustained swimming motility is required for maintaining intestinal spatial organization and persistence

Without swimming motility, *Vibrio* has clear defects in both immigration and intestinal persistence. Therefore, we sought to experimentally separate the roles motility plays during these different stages of colonization. We specifically wanted to determine whether *Vibrio* requires sustained motility for intestinal persistence or if the impaired immigration and altered assembly of motility mutant populations was in some way responsible for their aggregated and collapsing phenotype. To accomplish this, we built a motility “loss-of-function” switch that uses inducible CRISPR interference (CRISPRi) to suppress transcription of the flagellar motor gene operon *pomAB* (Figure 4A and Figure S3). The motility loss-of-function switch is based on a tetracycline induction system in which a constitutively expressed Tet repressor protein (TetR) is used to regulate the expression of a catalytically dead Cas9 (dCas9). We incorporated a constitutively expressed single guide RNA (sgRNA) to target dCas9 to the 5’ end of the native *pomAB* locus where it would block transcriptional elongation. To visually track switch activity in bacterial populations, we co-expressed *dcas9* with a gene encoding superfolder green fluorescent protein (sfGFP) (Figure 4A). To mark all cells independent of switch activity, we co-expressed a gene encoding dTomato with *tetR*. Details on switch design and optimization are provided in the Methods and in Figure S3A–S3D. We integrated the motility loss-of-function switch into the genome of wild-type *Vibrio* (creating *Vibrio*^motLOF^) and confirmed that induction of the switch with the tetracycline analog anhydrotetracycline (aTc) robustly inactivates swimming motility in vitro without perturbing growth (Figure S3E and S3F).

**Figure 4.**
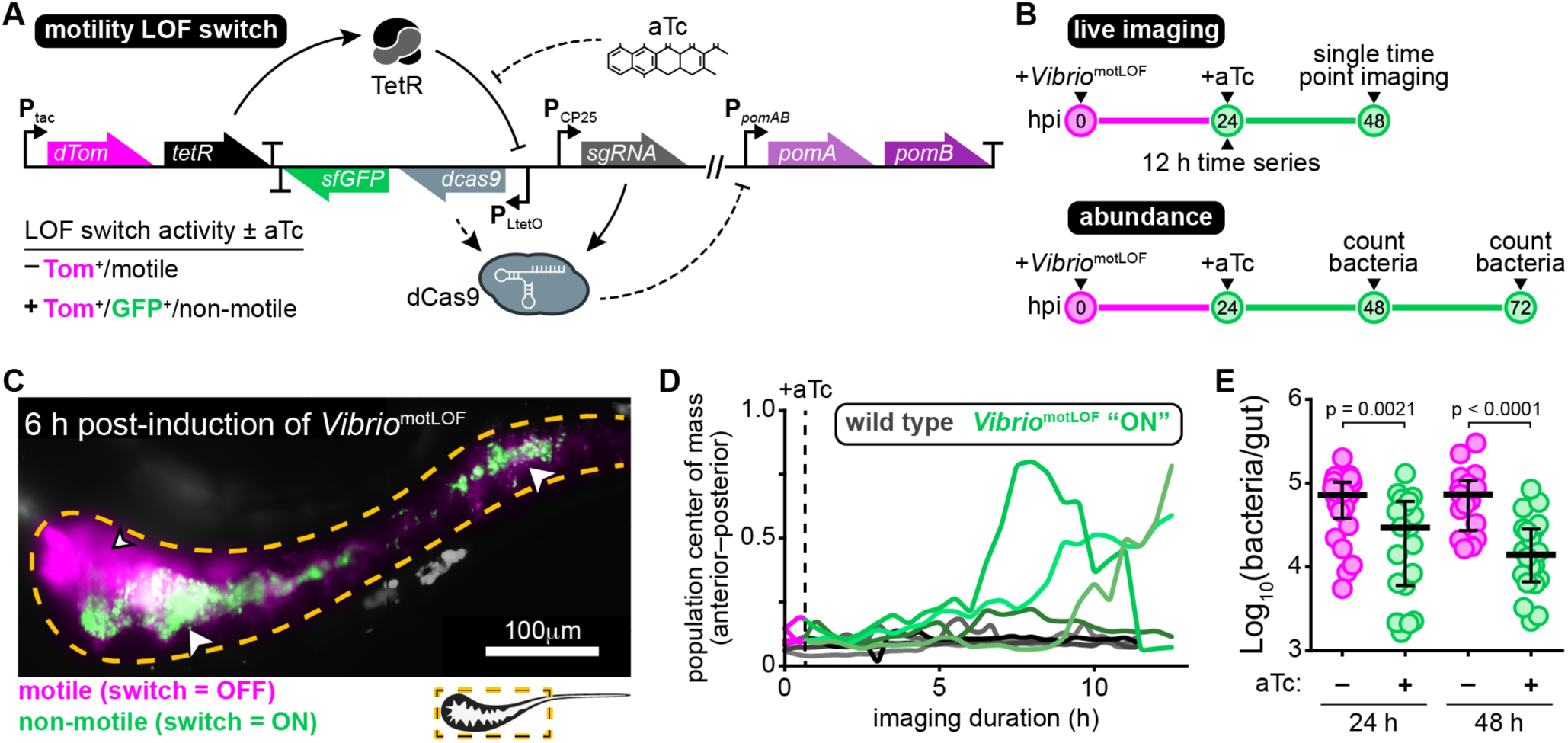
Sustained swimming motility is required for maintaining intestinal spatial organization and persistence. (see also Figure S3) **(A)** Schematic of CRISPRi-based motility loss-of-function (LOF) switch. Lower left table summarizes switch activity and bacterial behaviors plus/minus aTc. Bent arrows denote promoters, “T” denotes transcriptional terminators. Solid lines represent constitutive interactions, dashed lines represent induced interactions. **(B)** Experimental timelines used to investigate in situ inactivation of swimming motility. **(C)** A maximum intensity projection acquired by LSFM of an animal colonized by *Vibrio*^motLOF^ at 6 h post-induction. Dashed line mark approximate intestinal boundaries. An arrowhead with a black stroke marks an area of swimming cells expressing only dTomato (magenta, “switch = OFF”). White arrowheads mark aggregated cells (green, “switch = ON”). **(D)** Population center of mass over time for intestinal populations of wild-type *Vibrio* (gray) and *Vibrio*^motLOF^ (magenta/green). Lines are single bacterial populations within individual fish. Vertical dashed line marks time of aTc induction. **(E)** Abundances of *Vibrio*^motLOF^ at 24 and 48 h post-induction with aTc. Bars denote medians and interquartile ranges. Significant differences determined by Mann-Whitney.

With the motility loss-of-function switch constructed, we tested if sustained swimming motility is required by established *Vibrio* populations to persist within the intestine using both live imaging and cultivation-based measurements of abundance (Figure 4B). For live imaging, germ-free zebrafish were first colonized to carrying capacity with *Vibrio*^motLOF^. At 24 hpi, repression of motility was induced by adding aTc to the water of colonized zebrafish hosts. We then performed time series imaging of multiple animals using LSFM. Initially, subpopulations emerged that could be distinguished by their switch activation status, behavior, and spatial organization (Movie S5). Unswitched motile cells expressing only dTomato displayed a foregut localization pattern typical of wild-type *Vibrio* (Figure 4C). In contrast, we observed non-motile cells expressing GFP becoming aggregated and segregating away from motile populations (Figure 4C). GFP-positive cells within aggregates were more restricted to the intestinal lumen and their arrangement suggested they were encased in mucus (Figure 4C and Movie S5). By 10 hours post-induction, *Vibrio*^motLOF^ displayed clear shifts in population center of mass toward the midgut together with expulsion of multicellular aggregates (Figure 4D).

Cultivation-based measures of absolute abundances revealed that at 24 h post-induction *Vibrio*^motLOF^ populations had a ∼2.5-fold lower median abundance compared to uninduced controls (Figure 4E). Inducing for an additional 24 h resulted in a ∼5-fold reduction in median intestinal abundance (Figure 4E). Together, our experiments using the motility loss-of-function switch demonstrate that *Vibrio* requires sustained swimming motility to maintain its spatial organization and to persist at high levels. Our results also reveal that relatively brief interruptions in *Vibrio*’s swimming behavior are capable of producing rapid and dynamic changes in spatial organization and drops in abundance.

### Acquisition of swimming motility or chemotaxis leads to rapid recovery of intestinal spatial organization and abundance

We next asked whether established Δmot and Δche populations could recover their spatial organization and abundance if they reacquired swimming motility or chemotaxis, respectively. Answering this question would give insight into the capacity of resident gut bacteria and would-be pathobionts to exploit a sudden loss of host spatial control. Using the motility loss-of-function switch backbone, we constructed motility and chemotaxis “gain-of-function” switches by inserting either *pomAB* or *cheA2* in place of *dcas9* (Figure 5A). The motility and chemotaxis gain-of-function switches were integrated into the genomes of Δmot and Δche, respectively, creating Δmot^GOF^ and Δche^GOF^. In vitro tests showed that inducing the gain-of-function switches restored wild-type swimming behaviors in each strain without altering growth (Figure S4A–S4C). Moreover, activation of motility and chemotaxis prior to colonization produced intestinal abundances at 24 hpi that matched the carrying capacity of wild type (Figure 5B). These functional tests show that the motility and chemotaxis gain-of-function switches can be used to inducibly complement the Δmot and Δche mutants.

**Figure 5.**
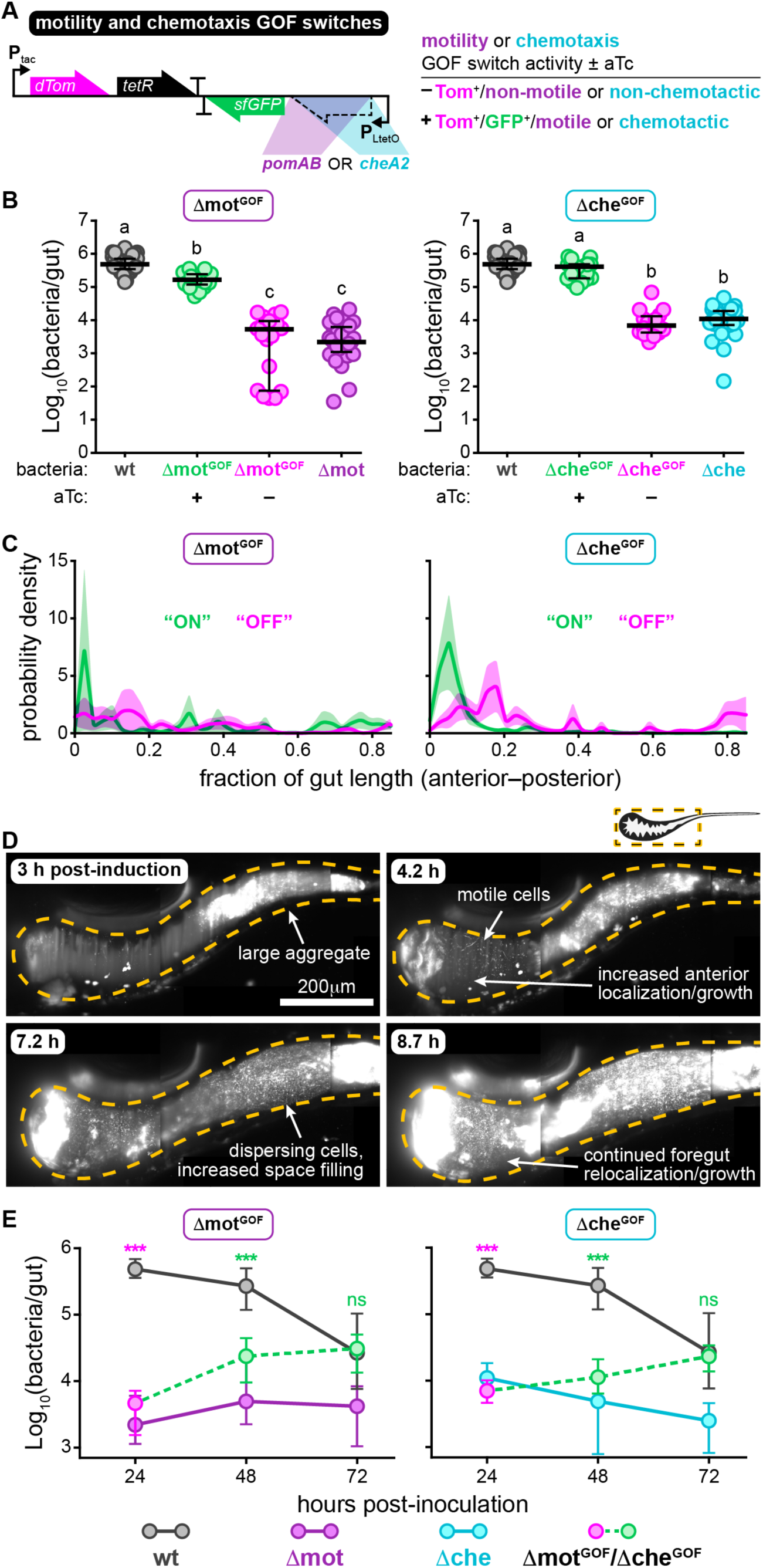
Acquisition of swimming motility or chemotaxis leads to rapid recovery of intestinal spatial organization and abundance. (see also Figure S4) **(A)** Schematic of the motility and chemotaxis gain-of-function (GOF) switches. Table summarizes switch activity and bacterial behaviors plus/minus aTc. **(B)** Δmot^GOF^ or Δche^GOF^ abundances 24 hpi plus/minus aTc. Δmot^GOF^ and Δche^GOF^ were pre-induced overnight in liquid culture prior to inoculation, aTc was maintained in the water for continuous switch activation. Abundances of wild-type *Vibrio*, Δmot, and Δche in wild-type hosts (from Figure 1A, 24 hpi) are shown for comparison. Bars denote medians and interquartile ranges. Letters denote significant differences. p < 0.05, Kruskal-Wallis and Dunn’s. **(C)** Probability densities showing the spatial distributions of Δmot^GOF^ and Δche^GOF^ at 24 h post-induction. Magenta = uninduced, green = induced. Shaded regions mark standard errors. Sample sizes (populations within individual animals): Δmot^GOF^ “OFF”, n = 5; Δmot^GOF^ “ON”, n = 7, Δche^GOF^ “OFF”, n = 6; Δche^GOF^ “ON”, n = 6. **(D)** Maximum intensity projections acquired by LSFM from the same animal showing Δche^GOF^ undergoing rapid changes in spatial organization following induction. Dashed lines mark approximate intestinal boundary. **(E)** Abundances of Δmot^GOF^ and Δche^GOF^ over time. Magenta and green circles indicate abundances plus/minus aTc, respectively. Plotted are medians and interquartile ranges (n ≥ 19 animals/marker). Abundances of wild-type *Vibrio*, Δmot, and Δche (from Figure 1A) are shown for comparison. Significant differences between each mutant and wild-type determined by Mann-Whitney (magenta asterisks: uninduced; green asterisks: induced). ***p < 0.0001, ns = not significant.

We monitored the response dynamics of activating swimming motility or chemotaxis in established populations following similar experimental timelines as depicted in Figure 4B. Live imaging revealed that induced populations of Δmot^GOF^ and Δche^GOF^ underwent clear shifts in spatial distribution toward the foregut within the first 24 h of induction compared to uninduced controls (Figure 5C). Strikingly, Δmot^GOF^ and Δche^GOF^ showed that large-scale changes in behavior and spatial organization could occur extremely rapidly, with both populations becoming more space-filling and foregut-localized within hours (Figure 5D, Movies S6 and S7). In contrast, cultivation-based measurements of absolute abundances showed only modest increases in median intestinal abundances in the first 24 h of induction (Figure 5E, 48 hpi). However, by 48 h post-induction the median intestinal abundances of Δmot^GOF^ and Δche^GOF^ populations had recovered to wild-type levels (Figure 5E, 72 hpi). Therefore, regaining swimming behavior and undergoing spatial reorganization precede the recovery of intestinal abundance.

Surprisingly, uninduced control populations of Δmot^GOF^ and Δche^GOF^ also exhibited a recovery in intestinal abundance by 72 hpi (Figure S4D). In vitro characterization and DNA sequencing revealed that this spontaneous recovery was likely due to non-synonymous mutations in *tetR* that were acquired during intestinal colonization and impaired the function of the Tet repressor protein, thus resulting in constitutive switch activation. While unexpected, we surmise that the extremely rapid sweep of “evolved clones” carrying disabled switches—which were rarely observed in induced populations or the aqueous environment outside the host (Figure S4E)—is evidence of strong selective pressures for motility traits within the gut.

### Motile bacterial cells within the intestine induce local and systemic *tnfa* expression

We next set out to connect *Vibrio*’s motility-based lifestyle to its pathobiont character. We recently showed that overgrowth of *Vibrio*-related taxa sparks intestinal pathology marked by increased epithelial hypertrophy and neutrophil influx that is dependent on TNFα signaling (Rolig et al., 2017). We further identified that *Vibrio* ZWU0020 on its own can potently stimulate inflammation (Rolig et al., 2015) and exacerbate pathology in susceptible hosts (Rolig et al., 2017). To explore the link between *Vibrio*’s motility behaviors and its inflammatory potential, we used LSFM and transgenic zebrafish hosts that express GFP under the control of the TNFα promoter (Tg(*tnfa*:GFP)) (Marjoram et al., 2015).

Germ-free animals displayed little *tnfa* reporter activity in or near the foregut where the bulk of wild-type *Vibrio* cells typically reside (Figure 6A). Similar to previous findings (Marjoram et al., 2015), animals colonized with a conventional, undefined microbial community also had low numbers of cells with *tnfa* reporter activity (Figure 6A). In contrast, at 24 hpi wild-type *Vibrio* induced pronounced *tnfa* reporter activity in numerous host cells within both the intestine and liver (Figure 6A). All animals colonized with wild-type *Vibrio* had *tnfa*-expressing cells in or near the liver whereas less than a third of germ-free and conventionalized animals had detectable *tnfa* reporter activity in this area (Figure 6B). Quantifying fluorescence intensity across the foregut region (including extraintestinal tissues and the liver) showed that *Vibrio* induces an approximately 100-fold increase in *tnfa* reporter activity over germ-free and conventional levels (Figure 6C). In contrast to wild-type *Vibrio*, Δmot and Δche elicited muted inflammatory responses. Animals colonized with Δmot showed a pattern of *tnfa* reporter activity similar to germ-free animals (Figure 6A–6C). However, despite the comparable intestinal abundances of Δmot and Δche (Figure 1A), Δche induced intermediate, although variable, levels of *tnfa* reporter activity (Figure 6A–6C). This finding suggests that host tissues do not merely sense bacterial abundances, but also their active swimming behavior and/or proximity to epithelial surfaces. Together, these data provide evidence that swimming motility and chemotaxis are major contributors to *Vibrio*’s proinflammatory potential.

**Figure 6.**
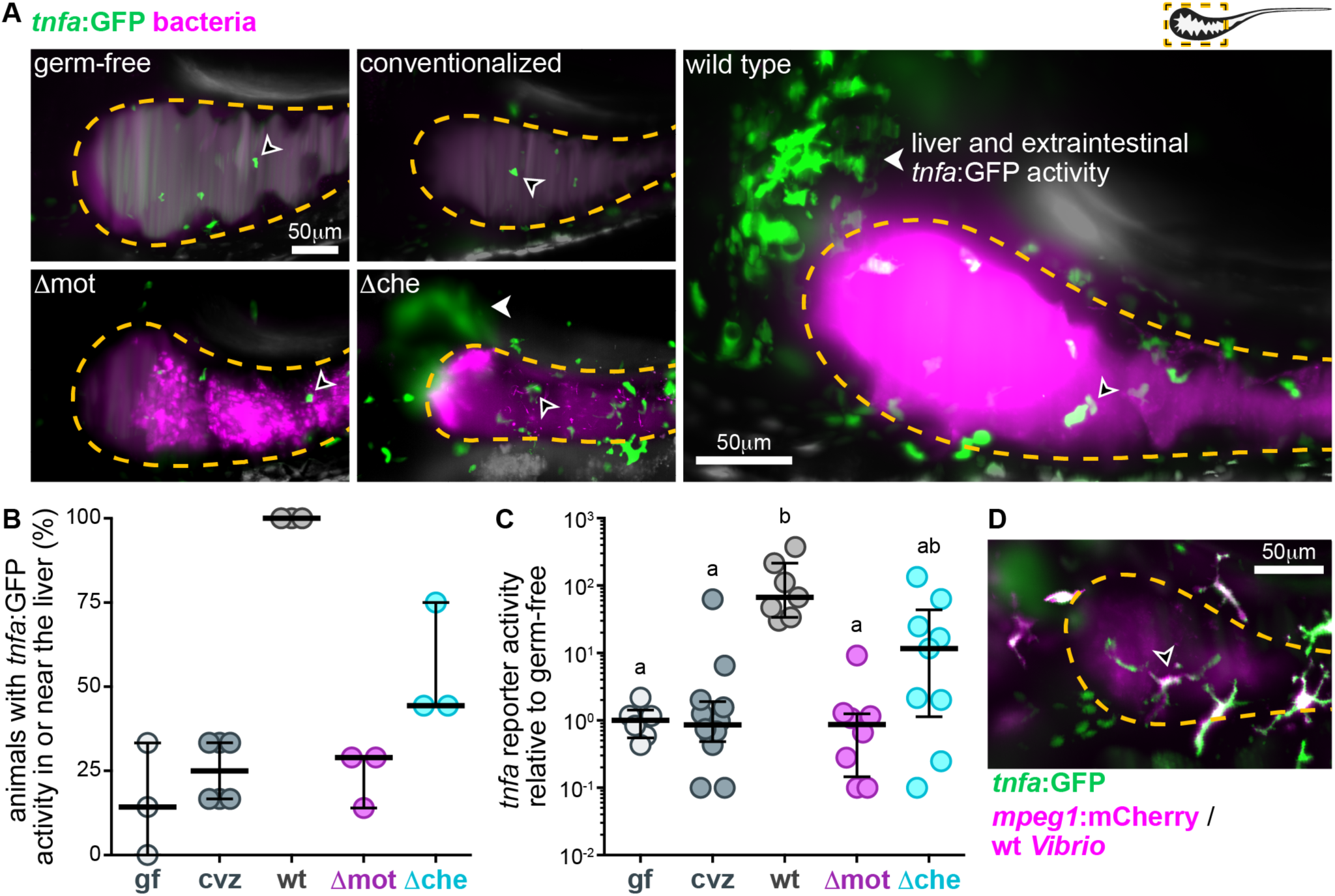
Motile bacterial cells induce local and systemic *tnfa* expression. **(A)** Maximum intensity projections acquired by LSFM of the foregut region of *tnfa*:GFP transgenic zebrafish raised germ-free, with a complex microbial community (conventionalized), or colonized solely with dTomato-expressing (magenta) wild-type *Vibrio*, Δmot, or Δche. Animals were imaged at 24 hpi. Dashed lines mark the approximate intestinal boundaries. Empty arrowheads mark host cells with *tnfa*:GFP reporter activity. Solid arrowheads mark *tnfa*:GFP reporter activity in extraintestinal tissues in or near the liver. **(B)** Percent of zebrafish subjected to different colonization regimes with *tnfa*:GFP activity in or near the liver. >6 animals/group were blindly scored by 3 researchers. Bars denote medians and interquartile ranges. gf: germ-free; cvz: conventionalized. **(C)** Total GFP fluorescence intensity across the foregut region normalized to median gf fluorescence intensity. Bars denote medians and interquartile ranges. Letters denote significant differences. p < 0.05, Kruskal-Wallis and Dunn’s. **(D)** Maximum intensity projections acquired by LSFM of the foregut region of a *tnfa*:GFP, *mpeg1*:mCherry (magenta) transgenic zebrafish colonized with dTomato-expressing wild-type *Vibrio* (magenta). Animal was imaged at 24 hpi. Open arrowhead indicates a *tnfa^+^*/*mpeg1^+^* cell.

We next probed the possible host cell types involved in sensing motile *Vibrio* populations. The amoeboid morphology and migratory behavior of many *tnfa*-expressing cells hinted that they might be immune cells (Figure 6A and Movie S8). Using double transgenic zebrafish carrying the *tnfa* reporter and expressing fluorescently marked macrophages (Tg(*mpeg1*:mCherry) (Ellett et al., 2011)), we found that ∼half (54 ± 10% [mean ± standard deviation, n = 100 cells from 4 animals]) of the *tnfa*-positive cells in the foregut region induced by *Vibrio* were indeed macrophages (Figure 6D and Movie S9). Nearly all *tnfa*-positive cells that were associated with intestinal tissues were macrophages (93 ± 12% [mean ± standard deviation, n = 18 cells from 3 animals]). In contrast, most *tnfa*-positive cells associated with the liver were not macrophages based on *mpeg1*:mCherry expression, nor were they neutrophils (based on experiments with animals carrying an *mpx*:mCherry reporter), suggesting that they were other non-immune cell types. Collectively, our data indicate that wild-type *Vibrio* populations stimulate expression of *tnfa* locally within intestinal tissues as well as at systemic sites, namely the liver. Macrophages are also one of the main cell types that is sensitive to *Vibrio* colonization.

### Host tissues rapidly respond to sudden increases in bacterial swimming motility within the intestine

To maintain homeostasis, the host must be simultaneously tolerant and sensitive to the activity of resident bacterial populations. It is crucial for host tissues to quickly differentiate between harmful and benign changes in the intestinal microbiota; for example, the overgrowth of a pathobiont versus diurnal fluctuations in commensal bacteria (Thaiss et al., 2016). Therefore, we next determined if sudden increases in bacterial motility behaviors—which are a potential signature of pathobionts escaping host control—could elicit an equally rapid host response.

Following a similar live imaging timeline as depicted in Figure 4B, we used LSFM to track *tnfa* reporter activity in response to induced populations of Δmot^GOF^. As expected, at time zero Δmot^GOF^ populations displayed low-abundance, high cohesion, and a posterior-shifted distribution with little *tnfa* reporter activity in host tissues (Figure 7A–7C). By 24 h post-induction, Δmot^GOF^ populations had begun to spatially reorganize within the foregut and contained an increased number of swimming cells (Figure 7A). At the same time, there was an increase in *tnfa*-expressing host cells near the intestine, which were likely macrophages (Figure 7A). In one instance, we captured *tnfa*-positive host cells within the mucosa adjacent to bacterial cells actively swimming near the epithelial surface (Movie S10). After the first 24 h of induction, the fraction of animals with *tnfa*-postive cells in or near the liver did not increase appreciably (Figure 7B); however, there was a ∼2.5-fold increase in median *tnfa* reporter activity, implying that initial responses to changes in bacterial swimming motility occur locally within intestinal tissues (Figure 7C). By 48 h of switch induction, Δmot^GOF^ populations exhibited wild-type-like space-filling properties and foregut-localization (Figure 7A). Likewise, *tnfa* reporter activity was also mostly restored to wild-type levels (Figure 7B and 7C). Nearly all animals (∼92%, n = 17) had *tnfa*-positive cells in or near the liver and the median *tnfa* reporter activity across the foregut region was 20-fold higher than germ-free levels (Figure 7B and 7C). Our data reveal that host tissues are remarkably sensitive to sudden increases in bacterial motility behaviors that occur over relatively short time scales.

**Figure 7.**
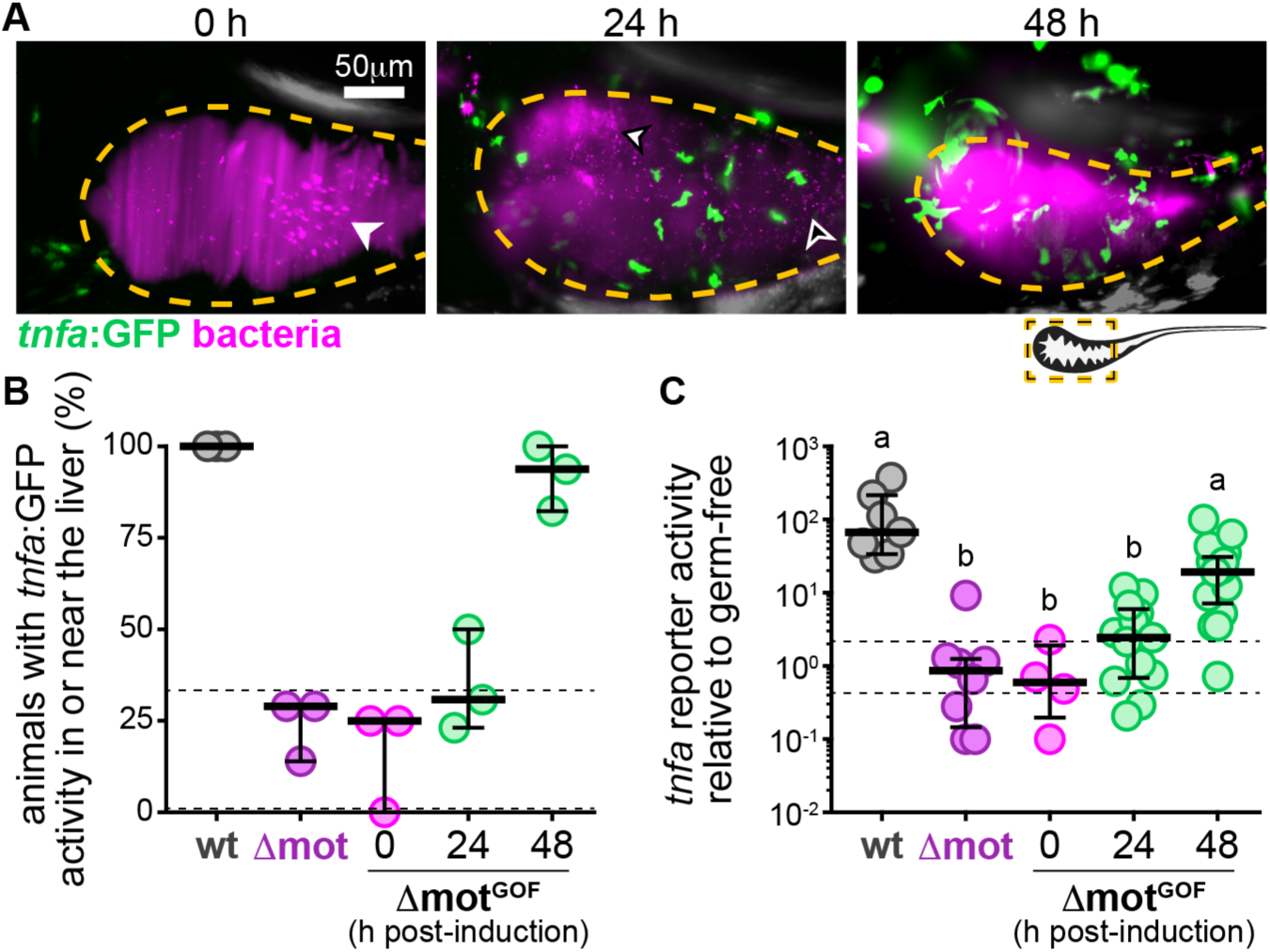
Host tissues rapidly respond to sudden increases in bacterial swimming motility within the intestine. **(A)** Maximum intensity projections acquired by LSFM of the foregut region of separate *tnfa*:GFP transgenic zebrafish colonized with Δmot^GOF^ (magenta). Dashed lines mark approximate intestinal boundaries. Times are h post-switch induction. Solid arrowhead marks bacterial aggregates, empty arrowhead marks single bacterial cells. **(B)** Percent of zebrafish subjected to different colonization regimes with *tnfa*:GFP activity in or near the liver. >4 animals/group were blindly scored by 3 researchers. Bars denote medians and interquartile ranges. Data from animals colonized with wild-type *Vibrio* (wt) or Δmot (from Figure 6B) are shown for comparison. Horizontal dashed lines mark gf range plotted in Figure 6B. **(C)** Total GFP fluorescence intensity across the foregut region normalized to median gf fluorescence intensity plotted in Figure 6C, horizontal dashed lines mark gf range. Bars denote medians and interquartile ranges. Data from animals colonized with wild-type *Vibrio* (wt) or Δmot (from Figure 6C) are shown for comparison. Letters denote significant differences. p < 0.05, Kruskal-Wallis and Dunn’s.

## DISCUSSION

Our study connects the motile lifestyle of a model pathobiont to its proinflammatory activity. All gut bacteria must contend with the intestine’s spatially dynamic landscape (Byndloss et al., 2018; Donaldson et al., 2015; Tropini et al., 2017). The mechanism by which *Vibrio* maintains stable colonization involves resisting intestinal flow through sustained swimming and chemotaxis. Conventional wisdom is that motility promotes the growth of bacteria by enabling them to forage nutrients and avoid hostile environments (Wei et al., 2011). In contrast, our data show that *Vibrio*’s motility behaviors within the zebrafish gut do not enhance its growth rate but rather allow it to counter intestinal flow. *Vibrio* thus provides a model of intestinal persistence that is distinct from more familiar examples involving adhesion to host tissues, which are largely based on examination of dissected and fixed samples (Donaldson et al., 2018; McLoughlin et al., 2016; Schluter et al., 2015). Ultimately, *Vibrio*’s colonization strategy uses continuous swimming to remain in place within the host’s intestine.

*Vibrio*’s swimming behavior underlies many of its pathobiont characteristics, including its ability to invade and displace resident bacteria, persist at high abundances, and stimulate host inflammation. The basis of *Vibrio*’s inflammatory activity is unknown, but future work is aimed at testing the contributions of its abundance, position along the intestine, mucosal proximity, and cellular behavior. Intriguingly, the different inflammatory activities of Δmot and Δche, despite their similar spatial organizations and production of flagella, highlights the possibility that the mechanism involves active bacterial motility. We posit that motility allows bacteria to access epithelial surfaces, increasing concentrations of inflammatory molecules at host cell surfaces and possibly triggering mechanosensory pathways. Flagellar rotation may also increase shedding of inflammatory components such as lipopolysaccharide and outer membrane vesicles, an idea supported by work in other *Vibrio* lineages (Aschtgen et al., 2016; Brennan et al., 2014).

*Vibrio*’s colonization dynamics show how swimming motility enables gut bacteria to evade spatial constraints imposed by the host. Notably, Δmot and Δche reveal how bacteria with impaired swimming motility surrender to intestinal mechanics, which entrap them within the lumen where they can be periodically purged. Thus, in addition to the intestine’s role in transporting digesta and expelling waste, intestinal flow and mucus dynamics appear to also exert spatial and population control over resident microbiota. Consistent with this, our characterization of *sox10* mutant zebrafish demonstrated how the enteric nervous system constrains intestinal bacterial abundances and composition, preventing dysbiosis (Rolig et al., 2017). From the bacterial side, we’ve also shown that sublethal antibiotics induce bacterial aggregation and enhance clearance from the intestine (Schlomann et al., 2019). A similar phenomenon was described in the mouse intestine, where antibody-mediated enchaining of bacterial cells enhanced clearance of *Salmonella* Typhimurium (Moor et al., 2017). Our observation that intestinal flow impacts the distribution of bacteria throughout the gut is also supported by findings in “gut-on-a-chip” fluidic systems (Cremer et al., 2016).

Given the clear advantage of motility behaviors within the gut, it is surprising that the majority of zebrafish gut bacteria studied so far—most of which are capable of flagellar motility— form aggregated populations made up of non-motile cells (Wiles et al., 2018). This discrepancy may be reconciled by considering the broader ecological life cycles of gut bacteria. For example, we have found that intestinal populations of *Aeromonas* grow more rapidly within multicellular aggregates than as planktonic cells (Jemielita et al., 2014), but *Aeromonas* also benefits from swimming motility for interhost dispersal (Robinson et al., 2018). *Aeromonas* thus highlights how aggregation and expulsion by intestinal flow may actually facilitate the transmission of resident gut bacteria and promote persistence within a population of hosts.

Further investigation of the relationship between intestinal mechanics and bacterial behaviors will open new avenues for therapeutic engineering of the gut microbiome. Our findings suggest that manipulating bacterial motility and aggregation may be used to induce large-scale, yet specific, changes in both bacterial abundances and host inflammatory state. Further, using drug- or diet-based modulators of intestinal flow may enhance the efficacy of antibiotics or promote microbiome recovery and fortification following perturbation. Our experiments using genetic switches to toggle bacterial motility and inflammatory activity serve as a proof-of-concept for these types of manipulations. Highlighting the potential of these interventions, human studies have shown that colonic transit time is a top predictor of microbiome composition (Falony et al., 2016; Roager et al., 2016). Moreover, impaired intestinal flow can lead to bacterial overgrowth and pathogenic changes in the microbiome (Dukowicz et al., 2007; Heanue and Pachnis, 2007; Rolig et al., 2017). Considering the dynamic nature of the intestinal ecosystem on spatial and temporal scales relevant to bacterial cells will be key to therapeutically engineering the microbiome.

## ACKNOWLEDGMENTS

Research was supported by an award from the Kavli Microbiome Ideas Challenge, a project led by the American Society for Microbiology in partnership with the American Chemical Society and the American Physical Society and supported by The Kavli Foundation. Work was also supported by the National Science Foundation under Awards 1427957 (R.P.) and 0922951 (R.P.), the M.J. Murdock Charitable Trust, and the National Institutes of Health (http://www.nih.gov/), under Awards P50GM09891 and P01GM125576 to K.G. and R.P., F32AI112094 to T.J.W., and T32GM007759 to B.H.S. R.B. was supported by NSF BIO/DBI Award 1460735 as visiting undergraduate intern. We also thank Rose Sockol and the University of Oregon Zebrafish Facility staff for fish husbandry, which are supported by a grant from the National Institute of Child Health and Human Development (P01HD22486). The funders had no role in study design, data collection and analysis, decision to publish, or preparation of the manuscript.

## AUTHOR CONTRIBUTIONS

Conceptualization, T.J.W., B.H.S., R.P., and K.G.; Methodology, T.J.W., E.S.W., and R.B.; Formal Analysis, T.J.W. and B.H.S.; Investigation, T.J.W., B.H.S., and E.S.W.; Writing – Original Draft, T.J.W. and B.H.S.; Writing – Review & Editing, T.J.W., B.H.S., E.S.W., R.P., and K.G.; Visualization, T.J.W., B.H.S.; Supervision, R.P., K.G.; Funding Acquisition, T.J.W., B.H.S., R.P., and K.G.

## STAR METHODS

### LEAD CONTACT AND MATERIALS AVAILABILITY

Further information and requests for resources and reagents should be directed to and will be fulfilled by the Lead Contact, Dr. Karen Guillemin.

(TBD) The following plasmids and associated sequences generated in this study will be deposited to Addgene:

1. pXS-GOF-switch
2. pXS-LOF-switch
3. pTn7xTS-GOF-switch
4. pTn7xTS-LOF-switch

### EXPERIMENTAL MODEL AND SUBJECT DETAILS

#### Animal care

All experiments with zebrafish were done in accordance with protocols approved by the University of Oregon Institutional Animal Care and Use Committee and following standard protocols (Westerfield, 2007). Specific handling and housing of animals during experiments are described in detail within the “METHODS DETAILS” section under the heading “Gnotobiology”. All zebrafish used in this study were larvae, between the ages of 4- and 7-days post-fertilization. Sex differentiation occurs later in zebrafish development and thus was not a factor in our experiments.

#### Zebrafish lines

Zebrafish lines used in this study included: University of Oregon stock wild-type ABCxTU; zebrafish carrying the *ret1^hu2846^* mutant allele (Ganz et al., 2018; Wiles et al., 2016); zebrafish carrying the Tg(*tnfa*:GFP) transgene (Marjoram et al., 2015); and zebrafish carrying the Tg(*mpeg1*:mCherry) transgene (Ellett et al., 2011). Double transgenic animals included Tg(*tnfa*:GFP) x Tg(*mpeg1*:mCherry) and Tg(*tnfa*:GFP) x Tg(*mpx*:mCherry) (Lam et al., 2013). Of note, *ret1^hu2846^* is recessive and adult zebrafish carrying this mutant allele were maintained as heterozygotes. Incrossing *ret1^hu2846^* animals produces *ret*^+/+^, *ret*^+/-^, and *ret*^-/-^ individuals. *ret*^-/-^ larvae can be visually distinguished from *ret*^+/+^ and *ret*^+/-^ larvae based on developmental features. In our study we classified *ret*^+/+^ and *ret*^+/-^ larvae together as “sibling” controls.

### METHOD DETAILS

#### Bacterial strains and culture

##### General

All wild-type and recombinant bacterial strains used or created in this study are listed in Table S1. Archived stocks of bacteria are maintained in 25% glycerol at −80°C. Prior to manipulations or experiments, bacteria were directly inoculated into 5 ml lysogeny broth (10 g/L NaCl, 5 g/L yeast extract, 12 g/L tryptone, 1 g/L glucose) and grown for ∼16 h (overnight) shaking at 30°C. For growth on solid media, tryptic soy agar was used. Gentamicin (10 μg/ml) was used to select recombinant *Vibrio* strains during their creation (for both gene deletion and insertion variants). Ampicillin (100 μg/ml) was used for maintaining plasmids in *E. coli* strains.

##### In vitro growth measurements

In vitro growth of bacterial strains was assessed using the FLUOstar Omega microplate reader. Prior to growth measurements, bacteria were grown overnight in 5 ml lysogeny broth at 30°C with shaking. The next day, cultures were diluted 1:100 into fresh lysogeny broth (aTc was added to the media when switch induction was required) and dispensed in triplicate or quadruplicate (i.e., 3–4 technical replicates) (200 μl/ well) into a sterile 96 well clear flat bottom tissue culture-treated microplate. Absorbance measurements at 600 nm were recorded every 30 min for 16 h (or until stationary phase) at 30°C with shaking. Growth measurements were repeated at least two independent times for each strain (i.e., two biological replicates) with consistent results. Data plotted are from a single replicate.

##### In vitro motility assays

The swimming behavior of each *Vibrio* strain was assessed using soft agar assays and live imaging of bacterial motility in liquid media on glass slides. For soft agar assays, bacteria were first grown overnight in 5 ml lysogeny broth at 30°C with shaking. One milliliter of bacterial culture was then washed by centrifuging cells at 7,000 x g for 2 minutes, aspirating media, and suspending in 1 ml 0.7% NaCl. This centrifugation and aspiration wash step was repeated once more and bacteria were suspended in a final volume of 1 ml 0.7% NaCl. One microliter of washed bacterial cells was inoculated into swim agar plates made of tryptic soy agar containing 0.2% agar. In the case of Δmot^GOF^ and Δche^GOF^, aTc was also added to the agar at the indicated concentrations. Swim plates were incubated at 30°C for 6 h and imaged using a Gel Doc XR+ Imaging System. For live imaging of swimming behavior, bacteria were first grown overnight in 5 ml lysogeny broth at 30°C with shaking. The next day, cultures of wild-type *Vibrio*, Δmot, and Δche were diluted 1:100 in tryptic soy broth and grown for 2 h with shaking at 30°C. Δmot^GOF^ and Δche^GOF^ were diluted 1:100 in tryptic soy broth +/– 50 ng/ml aTc and grown for 4 h with shaking at 30°C. *Vibrio*^motLOF^ was diluted 1:1000 in tryptic soy broth +/– 50 ng/ml aTc and grown for 7 h with shaking at 30°C. Prior to imaging, bacteria were diluted 1:40 in tryptic soy broth, mounted on glass slides with a coverslip and imaged for 10 s using a Nikon Eclipse Ti inverted microscope equipped with an Andor iXon3 888 camera. Representative maximum intensity projections of 10 s movies shown in Figures S1, S3, and S4 were generated in FIJI (Schindelin et al., 2012). For measurements of swimming behavior, bacteria were tracked using the radial center algorithm (Parthasarathy, 2012) for object localization and nearest-neighbor linking. Motility assays were repeated at least two independent times (i.e., two biological replicates) with consistent results.

##### Scanning electron microscopy

Bacteria were prepared for environmental scanning electron microscopy (ESEM) by first growing cells on tryptic soy agar overnight at 30°C. A sterile inoculating loop was used to transfer ∼100 μl of cells to a 1.6 ml tube containing 500 μl of 3% glutaraldehyde fixative. We visually confirmed that *Vibrio* cells isolated from an agar plate are highly motile and thus capable of producing flagella during culture on solid media. Cells were fixed overnight at 4°C. The next day, cells were sequentially washed in increasing concentrations of ethanol: first in plain ddH2O followed by 20%, 40%, 60%, and 80% ethanol. Each wash involved centrifuging cells at 7,000 x g for 2 minutes, aspirating media, and suspending in the next wash medium. A small aliquot of washed cell suspension was applied to a silicon wafer, dried, and imaged using a FEI Quanta 200 ESEM/VPSEM environmental scanning electron microscope provided by the University of Oregon’s Center for Advanced Materials Characterization in Oregon (CAMCOR) facility.

##### Disk diffusion assays

Disk diffusion assays were often used to test and optimize dTomato and sfGFP reporter function of genetic switches as described in Figure S3C and S3D. Bacteria were first grown overnight in 5 ml lysogeny broth at 30°C with shaking. One hundred microliters of dense overnight culture were spread onto tryptic soy agar plates to produce a lawn of growth. Prototyping was typically done using plasmid-base switches in *E. coli* (as was the case in Figure S3C and S3D), thus tryptic soy agar plates also contained ampicillin to ensure plasmid maintenance. A sterile piece of Whatman filter paper (∼0.5 cm wide) was placed in the center of the plate and impregnated with ∼2 μg of aTc. Plates were incubated overnight at 30°C. Switch reporter activity was assessed using a Leica MZ10 F fluorescence stereomicroscope equipped with 1.0x, 1.6x, and 2.0x objectives, and a Leica DFC365 FX camera. Images were captured and processed using standard Leica Application Suite software and FIJI (Schindelin et al., 2012).

#### Molecular techniques and genetic manipulations

##### General

Recombinant strains used or created in this study are listed in Table S1. Plasmids used or created in this study are listed in Table S2. Primer and oligo DNA sequences are listed in Table S3.

*E. coli* strains used for molecular cloning and conjugation were typically grown in 5 ml lysogeny broth at 30°C or 37°C with shaking in the presence of appropriate antibiotic selection to maintain plasmids. For propagation of *E. coli* on solid media, LB agar was used. Unless specified, standard molecular techniques were applied, and reagents were used according to manufacturer’s instructions. Restriction enzymes and other molecular biology reagents for polymerase chain reaction (PCR) and nucleic acid modifications were obtained from New England BioLabs. Various kits for plasmid and PCR amplicon purification were obtained from Zymo Research. The Promega Wizard Genomic DNA Purification Kit was used for isolating bacterial genomic DNA. DNA oligonucleotides were synthesized by Integrated DNA Technologies (IDT). Sanger sequencing was done by Sequetech to verify the sequence of all cloned genetic parts. A Leica MZ10 F fluorescence stereomicroscope with 1.0x, 1.6x, and 2.0x objectives and Leica DFC365 FX camera were used for screening fluorescent bacterial colonies.

Genome and gene sequences were retrieved from “The Integrated Microbial Genomes & Microbiome Samples” (IMG/M) website (https://img.jgi.doe.gov/m/) (Chen et al., 2017). Where applicable, “IMG” locus tags are provided for genetic loci, which can be used to access sequence information via the IMG/M website.

##### Construction of gene deletions

Markerless, in-frame gene deletions were constructed using allelic exchange and the pAX1 allelic exchange vector (Addgene Plasmid #117397) as previously described (Wiles et al., 2018). Detailed procedures and protocols can be accessed online: https://doi.org/10.6084/m9.figshare.7040264.v1. Creation of Δmot via deletion of *pomAB* (locus tags: ZWU0020_01568 and ZWU0020_01567) (Figure S1A) was reported previously (Wiles et al., 2018). Creation of Δche via deletion of *cheA2* (locus tag: ZWU0020_00514) (Figure S1A) was accomplished by first constructing a *cheA2* allelic exchange cassette using splice by overlap extension (SOE). The *cheA2* allelic exchange cassette was designed to fuse the start and stop codons of the *cheA2* gene (Figure S1A). PCR primer pairs WP165 + WP166 and WP167 + WP168 were used to amplify 5’ and 3’ homology regions flanking the *cheA2* gene, respectively, from *Vibrio* ZWU0020 genomic DNA. The resulting amplicons were spliced together and the SOE product was ligated into a pAX1-based allelic exchange vector, producing pAX1-ZWU0020-cheA2 (pTW383). After subsequent subcloning steps, the final sizes of the 5’ and 3’ homology regions were 763 bp and 845 bp.

The pAX1-ZWU0020-cheA2 vector was delivered into *Vibrio* via conjugation (i.e., bacterial mating) as previously described using *E. coli* SM10 as a donor strain (Wiles et al., 2018). Briefly, *Vibrio* and SM10/pAX1-ZWU0020-cheA2 were combined 1:1 on a filter disk placed on tryptic soy agar. The mating mixture was incubated at 30°C overnight. Following incubation, bacteria were recovered and spread onto tryptic soy agar containing gentamicin and incubated overnight at 37°C to select for *Vibrio* merodiploids. Merodiploid colonies were isolated and screened for successful deletion of the *cheA2* gene. Putative mutants were genotyped by PCR using primers that flanked the *cheA2* locus (WP0169 + CheA2.ZW20.KOconfirm.REV), which produced two differently sized amplicons representing the wild-type and mutant alleles (Figure S1A).

##### Design and construction of genetic switches

Customizable, plasmid-based gain-of-function (pXS-GOF-switch, pTW265) and loss-of-function (pXS-LOF-switch, pTW308) switch scaffolds were initially constructed and optimized using the pXS-dTomato (Addgene Plasmid #117387) backbone, which was previously generated (Wiles et al., 2018). The general architecture of switch elements is depicted in Figure S3A. Each element is flanked by unique restriction sites to allow straightforward insertion of new elements by restriction cloning. pXS-dTomato contains the “tracker” element, which comprises a constitutive Ptac promoter (without the lac operator sequence) (Wiles et al., 2018) driving the *dTomato* gene. The “switch reporter” element was first inserted, which comprises a PLtetO promoter (Lutz and Bujard, 1997) driving a *sfGFP* gene that was amplified from pTW168 using WP138 + WP118. Next, the “repressor” element was inserted, which comprises a *tetR* gene that was amplified from *Enterobacter* ZOR0014 genomic DNA using WP146 + WP139. As described in Figure S3B and S3C, a near-random ribosome binding site (ndrrdn) was incorporated by PCR into the 5’ untranslated region of the *tetR* gene via WP146. A clone containing the ribosome binding site sequence “ctaggt” was isolated that had strong reporter repression/induction and robust tracker expression. Next, as described in Figure S3B and S3D, a ribozyme-based insulator sequence (RiboJ) (Lou et al., 2012) was inserted between the switch reporter and the insertion site designated to hold switch “cargo” genes. The RiboJ sequence was inserted using a custom synthesized gBlock gene fragment (IDT). The resulting plasmid-based switch scaffold—comprising a tracker, switch reporter, repressor, and RiboJ sequence—became pXS-GOF-switch. To generate pXS-LOF-switch, we inserted the *dcas9* gene (Qi et al., 2013) (excised from pdCAS9, Addgene Plasmid #44249) as the “cargo” element and a constitutively expressed single guide RNA (“sgRNA” element) driven by the CP25 promoter (Jensen and Hammer, 1998), which was inserted using a custom synthesized gBlock gene fragment (IDT). The stock sgRNA that was inserted into the pXS-LOF-switch is based on a previously characterized sgRNA specific for the *lacZ* gene of *E. coli* (Qi et al., 2013), which facilitated optimization of loss-of-function switch activity in *E. coli* K-12 (MG1655). To expedite insertion of the gain-of-function and loss-of-function switches into the *Vibrio* chromosome, each switch scaffold was subcloned into the previously described Tn*7* delivery vector pTn7xTS (Addgene Plasmid #117389), creating pTn7xTS-GOF-switch (pTW285) and pTn7xTS-LOF-switch (pTW317). We note that insertion of the switch scaffolds into the pTn7xTS vector limits some downstream customization due to restriction site conflicts.

To construct the motility loss-of-function switch, the *lacZ* sgRNA in the pTn7xTS-LOF-switch was replaced with a sgRNA specific for the *Vibrio pomA* gene, creating pTn7xTS-mot-LOF-switch (pTW340). The *pomA* sgRNA was inserted using a custom synthesized gBlock gene fragment (IDT). To construct the motility gain-of-function switch, the *pomAB* locus, including the native *pomA* ribosome binding site, was amplified using WP170 + WP171 and inserted into the cargo site of pTn7xTS-GOF-switch, creating pTn7xTS-mot-GOF-switch (pTW324). To construct the chemotaxis gain-of-function switch, the *cheA2* locus, including the native *cheA2* ribosome binding site, was amplified using WP92 + WP93 and inserted into the cargo site of pTn7xTS-GOF-switch, creating pTn*7*xTS-che-GOF-switch (pTW282).

##### Tn7-mediated chromosomal insertions

Chromosomal insertion of fluorescent markers and genetic switches was done via a Tn*7* transposon-based approach using the Tn*7* delivery vector pTn7xTS as previously described (Wiles et al., 2018). Detailed procedures and protocols can be accessed online: https://doi.org/10.6084/m9.figshare.7040258.v1. Specific pTn7xTS vectors carrying markers or switches were delivered into *Vibrio* via triparental mating using two *E. coli* SM10 donor strains carrying either the pTn7xTS delivery vector or the pTNS2 helper plasmid (Addgene Plasmid #64968). Briefly, *Vibrio* and SM10 donor strains were combined 1:1:1 on a filter disk placed on tryptic soy agar. The mating mixture was incubated at 30°C overnight. Following incubation, bacteria were recovered and spread onto tryptic soy agar containing gentamicin and incubated overnight at 37°C to select for *Vibrio* insertion variants. Insertion of the Tn*7* transposon and the genetic cargo it carried into the *attTn7* site near the *glmS* locus of *Vibrio* was confirmed by PCR using primers WP11 + WP12.

Fluorescently marked wild-type *Vibrio* constitutively expressing dTomato (ZWU0020 *attTn7*::*dTomato*) was previously generated using pTn7xTS-dTomato (Addgene Plasmid #117391) (Wiles et al., 2018). In the current work, fluorescenlty marked Δmot and Δche were constructed in the same way, creating Δmot *attTn7*::*dTomato* and Δche *attTn7*::*dTomato*. *Vibrio*^motLOF^ was created by inserting the motility loss-of-function switch from pTn7xTS-mot-LOF-switch. Δmot^GOF^ was created by inserting the motility gain-of-function switch from pTn7xTS-mot-GOF-switch. Δche^GOF^ was created by inserting the chemotaxis gain-of-function switch from pTn7xTS-che-GOF-switch.

#### Gnotobiology

##### Germ-free derivation

For all experiments, zebrafish embryos were initially derived germ-free using previously described gnotobiotic procedures with slight modification (Melancon et al., 2017). Briefly, fertilized eggs from adult mating pairs were harvested and incubated in sterile embryo media (EM) containing ampicillin (100 μg/ml), gentamicin (10 μg/ml), amphotericin B (250 ng/ml), tetracycline (1 μg/ml), and chloramphenicol (1 μg/ml) for ∼6 h. Embryos were then washed in EM containing 0.1% polyvinylpyrrolidone–iodine followed by EM containing 0.003% sodium hypochlorite. Surface sterilized embryos were distributed into T25 tissue culture flasks containing 15 ml sterile EM at a density of one embryo per milliliter and kept in a temperature-controlled room at 28–30°C with a 14 h/ 10 h light/dark cycle. The germ-free status of larval zebrafish was assessed before every experiment by visually inspecting flask water for microbial contaminants using an inverted microscope. Culture-based assessment of germ-free status was done as needed by plating 100 μl flask water on rich media (e.g., tryptic soy agar). Embryos were sustained on yolk-derived nutrients and not fed prior to or during any experiments.

##### Bacterial associations

For bacterial associations, bacterial strains were grown overnight in lysogeny broth with shaking at 30°C and prepared for inoculation by pelleting the cells from 1 ml of culture for 2 min at 7,000 x g and washed once in sterile EM. For all experiments, except where noted otherwise, washed bacteria were inoculated into the water of T25 flasks containing 4-day-old larval zebrafish at a final density of ∼10^6^ bacteria/ml. For competition experiments, *Vibrio* strains were added to the water of *Aeromonas*-colonized zebrafish (at 5-days-old) without removing the original *Aeromonas* inoculum from the water. In addition, to enable enumeration of *Aeromonas* and *Vibrio* strains on agar plates, competition experiments were done using a previously constructed dTomato-expressing *Aeromonas* strain (*Aeromonas attTn7*::*dTomato*) (Wiles et al., 2018). For loss-of-function and gain-of-function switch experiments involving cultivation-based abundance measurements, prior to aTc-induction zebrafish were washed and placed in sterile EM to ensure that changes in intestinal populations were not interfered with by bacteria in the water. To conventionalize animals (i.e., colonize with a complex, undefined microbial consortium), 0 and 4-day-old larval zebrafish were inoculated with 100 μl of water taken from parental spawning tanks. No difference was found between conventionalization times in terms of host *tnfa*:GFP expression.

#### Cultivation-based measurement of abundances

Dissection of larval zebrafish guts was done as previously described with slight modification (Milligan-Myhre et al., 2011). Briefly, dissected guts of tricaine-euthanized zebrafish were harvested and placed in a 1.6 ml tube containing 500 μl sterile 0.7% saline and 100 μl 0.5 mm zirconium oxide beads. Guts were homogenized using a bullet blender tissue homogenizer for 25 seconds on power 4. Lysates were serially plated on tryptic soy agar and incubated overnight at 30°C prior to enumeration of colony forming units and determination of bacterial abundances. Abundance data presented throughout the main text and in Figure S2 are pooled from a minimum of two independent experiments (n = 16–36 dissected guts per condition). Abundance data presented for Δmot^GOF^ and Δche^GOF^ without aTc induction in Figure S4 are from a single representative experiment (n = 8–10 dissected guts per condition; water abundances are from single measurements). Samples with zero countable colonies on the lowest dilution were set to the limit of detection (5 bacteria per gut). Data were plotted and analyzed using GraphPad Prism 6 software. Unless stated otherwise, statistical differences between two groups of data were determined by Mann-Whitney; statistical differences between two paired groups of data were determined by Wilcoxon; and statistical differences among three or more groups of data were determined by Kruskal-Wallis test with Dunn’s multiple comparisons test.

#### Live imaging

##### Light sheet fluorescence microscopy

Live larval zebrafish were imaged using a custom-built light sheet fluorescence microscope previously described in detail (Jemielita et al., 2014). Prior to mounting, larvae were anesthetized with MS-222 (tricaine). A metal plunger was used to mount fish into small glass capillaries containing 0.5% agarose gel. Samples were then suspended vertically, head up, in a custom imaging chamber containing embryo media and anesthetic. Larvae in the set gel were extruded from the end of the capillary and oriented such that the fish’s left side faces the imaging objective. For experiments involving just fluorescent bacteria, the ∼ 1mm long intestine is imaged in four subregions that are registered in software after imaging. A single 3D image of the full intestine volume (∼200×200×1200 microns) sampled at 1-micron steps between z-planes is imaged in ∼45 seconds. For experiments including the *tnfa*:GFP reporter, only one subregion containing the anterior foregut region and ∼100 microns of tissue anterior to the gut was captured for the majority of samples. In these image stacks, nearly the full extent of the fish’s left-right width was captured, approximately 400 microns in z. For time lapse imaging of genetic switch induction, fish were mounted as normal and baseline dynamics were captured for 30-90 min depending on the experiment. Then, the inducer aTc was added to the sample chamber media in an approximately 1 ml solution of embryo media, MS-222, and aTc. Excitation lasers of wavelengths 488 and 561 nm were adjusted to a power of 5 mW as measured before the imaging chamber. An exposure time of 30 ms was used for all 3D scans and 2D movies. Time lapse imaging was performed overnight, except for the additional growth rate measurement for Δche, which occurred during the day. For color images presented within figures, autofluorescent tissues— namely, the yolk, swim bladder, and ventral skin—were manually converted to grayscale to enhance clarity.

##### Identification of fluorescent bacteria

Identification of bacteria in zebrafish images was conducted using a previously described computation pipeline written in MATLAB (Jemielita et al., 2014; Schlomann et al., 2018). In brief, individual bacteria are first identified with a combination of wavelet filtering (Olivo-Marin, 2002), standard difference of Gaussians filtering, intensity thresholding, and manual curation. Then, multicellular aggregates, which are too dense to resolve individual cells, are segmented via a graph cut algorithm (Boykov and Kolmogorov, 2004) seeded with an intensity mask. The number of cells per multicellular aggregate is estimated by dividing the total aggregate fluorescence intensity by the mean intensity of single cells. These estimates of number of cells per bacterial object in the gut are then used to compute spatial distributions along the length of the gut, following a manually drawn line drawn that defines the gut’s center axis.

##### Measurement of in vivo growth rates

Through time-lapse imaging and the computational image analysis methods discussed above, bacterial growth rates in the intestine can be directly measured by linear fits to log-transformed abundances (Jemielita et al., 2014; Schlomann et al., 2019; Wiles et al., 2016). The in vivo growth rate of wild-type *Vibrio* was previously measured (Wiles et al., 2016). The in vivo growth rate for Δmot was measured in the time traces shown in Figure 3A, using manually defined windows of clear exponential growth. To exclude effects of density dependence on the growth rate, only those traces that began at least 1 order of magnitude below the median Δmot abundance were considered. For Δche, the time traces in Figure 3A were insufficient for growth rate estimation, because abundances remained at high levels for most of the experiment. Therefore, we measured the growth rate in populations shortly after initial colonization. Specifically, Δche was allowed to colonize germ-free fish for 6 hours, after which fish were mounted for time lapse imaging. Previous work on another zebrafish gut bacterial symbiont showed that exponential growth rates in established and nascent populations are equal (Wiles et al., 2016). Abundance data for these time traces are included in the Data S1.

##### Quantification of tnfa:GFP fluorescence

Cells and tissues expressing *tnfa*:GFP were segmented in 3D with basic intensity threshold-based segmentation. A pixel intensity threshold of 1500 was empirically found to be a conservative threshold and was used for all samples. The 488 nm excitation laser power was set at 5 mW prior to entering the sample chamber for all samples. The camera was a pco.edge scientific CMOS camera (PCO, Kelheim, Germany). The resulting identified objects were then filtered by size to remove noise. GFP signal from near the ventral skin was excluded with a manually defined cropped region created in the ImageJ software (Schindelin et al., 2012). Green autofluorescence from the interior gut region rarely passed the intensity and size thresholds to contribute to measured *tnfa*:GFP signal. Similarly, in the motility gain-of-function switch experiments, we found that the GFP reporter of switch induction never reached fluorescence intensity levels high enough to contribute measurably to the *tnfa*:GFP signal. Nevertheless, this region was automatically identified and removed via intensity threshold-based segmentation in the red 568/620 nm (excitation/emission) color channel. Both red autofluorescence and signal from red (dTomato) fluorescent bacteria were used to identify this gut region. Finally, to standardize total *tnfa*:GFP quantification across different samples, an operational peri-intestinal region was defined as containing the foregut plus all tissue 100 microns anterior of the start of the gut, which was automatically identified in the mask generated from the red color channel. Automatic gut segmentation and removal was not performed for the dual *tnfa*:GFP/*mpeg1*:mCherry reporter fish.

##### Measuring tnfa:GFP/mpeg1:mCherry fluorescence

Red fluorescence from *mpeg1*:mCherry marking macrophages was segmented analogously to *tnfa*:GFP signal, using basic intensity threshold-based segmentation in 3D and size filtering. A *tnfa*+ object and an *mpeg*+ object were considered to overlap if their centroids were separated by less than 10 microns, a threshold empirically determined to produce accurate results as judged by eye. The fraction of *tnfa*+ objects that were also *mpeg*+ and the fraction of *mpeg*+ objects that were also *tnfa*+ were computed using the counts for overlapping and non-overlapping cells.

## QUANTIFICATION AND STATISTICAL ANALYSIS

Data were plotted using MATLAB and GraphPad Prism 6 software. Statistical analyses were done using GraphPad Prism 6. Unless stated otherwise, medians and interquartile ranges were plotted. Statistical tests performed are specified in figure legends and within the Methods under “Cultivation-based measurement of abundances”. A p-value of 0.05 or less was considered significant for all analyses. Sample sizes are noted within the main text, figure legends, within the Methods under “Cultivation-based measurement of abundances”, and in Data S1 (.xls). What “n” represents is specified in the main text and figure legends.

## DATA AND CODE AVAILABILITY

All numerical data generated or analyzed during this study are provided in Data S1 (.xls). All code used in this study was based on previously published algorithms and is available upon request. Size and volume of raw imaging data prevents uploading to a public repository but will be made available upon request.

## MOVIE LEGENDS

**Movie S1. Montage of real time movies showing wild-type *Vibrio*, Δmot, and Δche within larval zebrafish intestines**. (associated with Figure 2)

Movies were acquired by light sheet fluorescence microscopy at 24 hpi. Wild-type *Vibrio* is highly motile and planktonic, with swimming cells frequently making close contact with the intestinal epithelium. The bright signal in the left side of the frame is a mass of motile cells that is too dense for individuals to be resolved (see Figure 2C). In contrast, Δmot is largely aggregated and confined to the lumen. The Δche mutant exhibits an intermediate phenotype consisting of a motile subpopulation that is less dense than wild-type populations. The field of view centers on the foregut region. Scale bar = 50 μm.

**Movie S2. Montage of animated z-stacks showing wild-type *Vibrio*, Δmot, and Δche within larval zebrafish intestines**. (associated with Figure 2)

Movies were acquired by light sheet fluorescence microscopy at 24 hpi. Wild-type *Vibrio* is highly motile and planktonic, with swimming cells frequently making close contact with the intestinal epithelium. The bright signal in the left side of the frame is a mass of motile cells that is too dense for individuals to be resolved (see Figure 2C). In contrast, Δmot is largely aggregated and confined to the lumen. The Δche mutant exhibits an intermediate phenotype consisting of a motile subpopulation that is less dense than wild-type populations. The field of view centers on the foregut region. The label in the upper-right corner denotes the depth in z (left-right) through the intestine. Scale bar = 50 μm.

**Movie S3. Montage of time-lapse movies showing wild-type *Vibrio*, Δmot, and Δche within larval zebrafish intestines**. (associated with Figure 3)

Movies were acquired by light sheet fluorescence microscopy starting at ∼24 hpi. Wild-type *Vibrio* cells, which are highly motile and planktonic, robustly localize to the foregut region. The bright signal in the left side of the frame is a stable mass of motile cells that is too dense for individuals to be resolved (see Figure 2C). In contrast, Δmot is largely aggregated, confined to the lumen, and exhibits large fluctuations in spatial organization, including the rapid expulsion of a large aggregate. The Δche mutant exhibits an intermediate phenotype, consisting of a motile subpopulation that is less dense than wild-type populations with large, multicellular aggregates. A large aggregate of Δche cells is expelled near the end of the movie. The field of view spans the entire larval intestine. Scale bar = 200 μm.

**Movie S4. Animation of the spatiotemporal dynamics of wild-type *Vibrio*, Δmot, and Δche within larval zebrafish intestines**. (associated with Figure 3)

Through computational image analysis, bacterial populations were segmented and enumerated. From this quantification, we computed the fraction of the population that were single cells (planktonic fraction) and computed the population center of mass along the length of the gut (population center). Each marker represents an entire bacterial population from an individual fish. The movie depicts the time evolution of multiple populations in this 2-dimensional phase space. Wild-type *Vibrio* populations robustly localize to the foregut region and maintain a high planktonic fraction. In contrast, Δmot and Δche populations undergo large fluctuations in aggregation and localization over time.

**Movie S5. Inactivation of swimming motility in established *Vibrio*^motLOF^ populations using the motility loss-of-function switch**. (associated with Figure 4)

Shown are two examples of *Vibrio*^motLOF^ switching dynamics within the larval zebrafish intestine captured by light sheet fluorescence microscopy. *Vibrio*^motLOF^ initially colonized each intestine in a phenotypically wild-type state (i.e., switch = “OFF”) with cells expressing only dTomato (magenta) and displaying a strong localization to the foregut and a high fraction of motile cells. At time zero, populations were induced by addition of aTc to the media. Both examples show the emergence of a multicellular aggregate from the anterior population of motile cells, a posterior shift in overall distribution, and an increase in GFP expression signaling switch activation. Scale bars for time-lapses are 200 μm. Each frame of the time-lapses are maximum intensity projections of a 3D image stack across the full intestinal volume. For the second example, we highlight the 3D structure of an emerging bacterial aggregate (arrow) with an animated rendering (dTomato fluorescence only). Scale bar for the rendering is 50 μm. The montage ends with a real time movie of *Vibrio*^motLOF^ cells approximately 16 h post-induction showing widespread loss of motility (dTomato fluorescence only). Real time movie scale bar = 50 μm.

**Movie S6. Activation of swimming motility in an established Δmot^GOF^ population using the motility gain-of-function switch**. (associated with Figure 5)

Δmot^GOF^ initially colonized the intestine with the motility gain-of-function switch in the “OFF” state and therefore was non-motile and assembled a population that was aggregated and had a poster-shifted distribution. At time zero, the population was induced by addition of aTc to the media. The resulting switching dynamics were captured by light sheet fluorescence microscopy. Each frame of the time-lapse is a maximum intensity projection of a 3D image stack across the full intestinal volume. Over time, motile cells appear and occupy the foregut region. Scale bar = 200 μm. Following the time-lapse, we show a real time movie of a different fish at approximately 6 h post-induction that captures induced Δmot^GOF^ cells swimming within the foregut. Real time movie scale bar = 50 μm.

**Movie S7. Activation of chemotaxis in an established Δche^GOF^ population using the chemotaxis gain-of-function switch.** (associated with Figure 5)

Δche^GOF^ initially colonized the gut with the chemotaxis gain-of-function switch in the “OFF” state and therefore was non-chemotactic and assembled a population that was aggregated and had a poster-shifted distribution. At time zero, the population was induced by addition of aTc to the media. The resulting switching dynamics were captured by light sheet fluorescence microscopy. Each frame of the time-lapse is a maximum intensity projection of a 3D image stack across the full intestinal volume. Over time, there is a dramatic increase in the number of planktonic and motile cells that occupy the foregut region. Scale bar = 200 μm. Following the time-lapse, we show a real time movie of a different fish at approximately 6 h post-induction that captures induced Δche^GOF^ cells swimming within the foregut. Real time movie scale bar = 50 μm.

**Movie S8. Migratory behavior of *tnfa*:GFP^+^ cells.** (associated with Figure 6)

Time-lapse movie of a live *tnfa*:GFP transgenic larval zebrafish showing the migratory behavior of gut-associated *tnfa*^+^ cells (arrowheads). Images were acquired by light sheet fluorescence microscopy. Each frame of the time-lapse is a maximum intensity projection of a 3D image stack that captures the full intestinal volume. Scale bar = 200 μm.

**Movie S9. Animated z-stack of a *tnfa*:GFP/*mpeg1*:mCherry double transgenic larval zebrafish colonized with wild-type *Vibrio*.** (associated with Figure 6)

The *mpeg1*:mCherry reporter and *Vibrio* dTomato marker were imaged simultaneously using a single excitation and emission system, and are shown in magenta. *tnfa*:GFP fluorescence is shown in green. Images were acquired by light sheet fluorescence microscopy. We first show an animated z-stack that depicts single planes of the light sheet with the depth (left–right) indicated in the upper right. *tnfa*^+^ and *mpeg1*^+^ single-positive cells, as well as *tnfa*^+^/*mpeg1*^+^ double-positive cells, are apparent. Scale bar = 50 μm. Following the animated z-stack, we show a two-color, 3D rendering. Rendering scale bar = 50 μm.

**Movie S10. Spatial distribution of *tnfa*^+^ host cells responding to swimming bacterial cells within the intestine.** (associated with Figure 7)

Montage shows the foregut region of a larval zebrafish carrying the *tnfa*:GFP reporter (green) colonized with Δmot^GOF^ (magenta) 24 h post-induction of the motility gain-of-function switch with aTc. Images and real time movie were acquired by light sheet fluorescence microscopy. We first show an animated z-stack that depicts single planes of the light sheet with the depth (left–right) indicated in the upper right. Arrows indicate *tnfa*^+^ host cells and bacterial cells. Next, we show a 3D rendering of the same intestine, which highlights the association of *tnfa*^+^ host cells with the mucosa. The montage ends with a real time movie of a single optical plane showing the swimming behavior of induced Δmot^GOF^ cells relative to *tnfa*^+^ host cells within the mucosa. All scale bars = 50 μm.

## EXCEL TABLE

**Data S1.** File (.xls) containing all plotted numerical data.

## SUPPLEMENTAL FIGURES & TABLES

**Figure S1.**
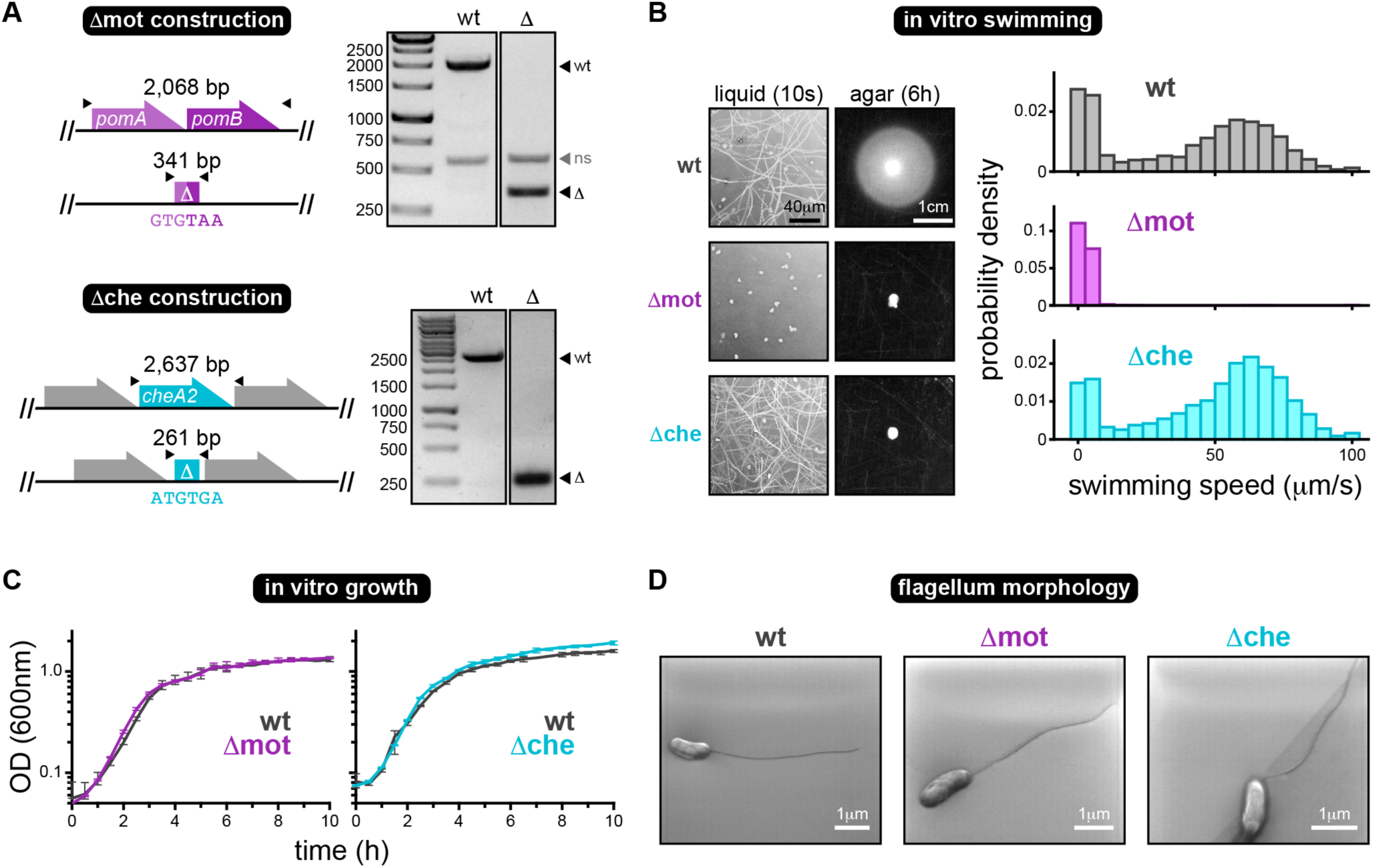
Motility and chemotaxis mutant construction and in vitro characterization. (associated with Figures 1 and 2) **(A)** Gene diagrams depict the in-frame markerless deletion of *pomAB* (Δmot construction) and *cheA2* (Δche construction). “Δ” denotes the mutant allele and the DNA sequence shown below represents the resulting fusion of the start and stop codons in each case. Black triangles represent primers used for PCR-confirmation of each mutant, and the amplicon sizes (bp) of the wild-type (wt) and mutant (Δ) alleles are provided above each locus. DNA gels to the right of each diagram show the successful deletion of both *pomAB* and *cheA2* from the *Vibrio* chromosome. We note that the Δmot mutant was constructed in a previous publication (Wiles et al., 2018) and the DNA gel shown is a version of that already published but is included here for continuity and thoroughness. ns: non-specific amplicon. **(B)** Left: Swimming motility of wild type (wt), Δmot, and Δche in liquid media and soft agar. Motility in liquid media was recorded for 10 seconds on a glass slide. Images show cellular movements over the entire 10 second period, which illustrates each cell’s swimming trajectory. Swim distances were captured 6 h post-inoculation of bacteria into the agar. Right: Probability densities showing the distribution of cellular swimming speeds in liquid media for each *Vibrio* strain. Sample sizes (measured bacterial swim tracks): wt = 2,962; Δmot = 754; Δche = 3,069. **(C)** In vitro growth curves of each *Vibrio* strain in rich media (lysogeny broth). Line traces the average optical density (OD) from four replicate wells, bars indicate range. **(D)** Scanning electron micrographs of each *Vibrio* strain after growth on solid media. Images show that each strain is capable of assembling a single polar flagellum.

**Figure S2.**
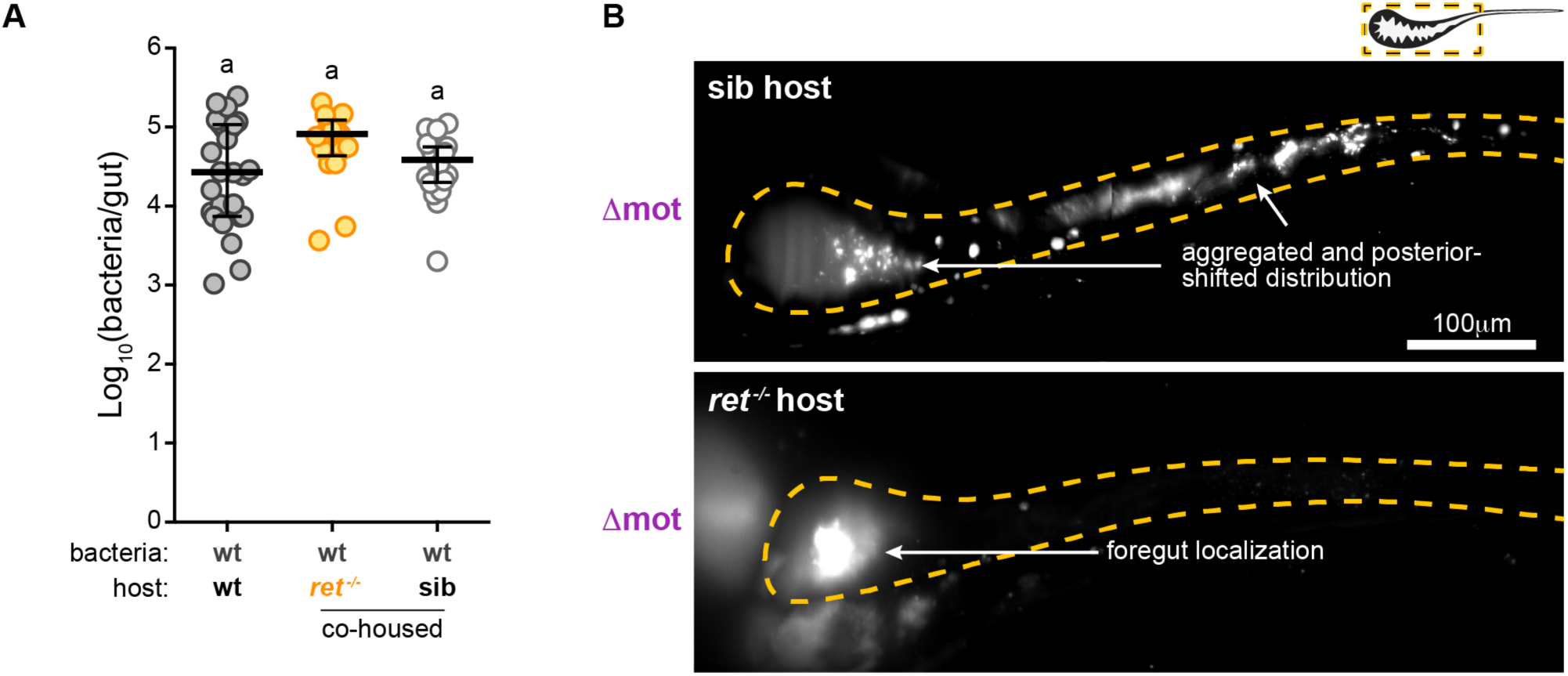
Additional wild-type and Δmot colonization data in *ret*^-/-^ mutant hosts. (associated with Figure 3) **(A)** Cultivation-based abundances for wild-type *Vibrio* in co-housed *ret*^-/-^ mutant hosts and wild-type/heterozygous sibling controls (sib). Abundances of wild-type *Vibrio* in wild-type hosts (from Figure 1A, 72 hpi) are shown for comparison. Letters denote significant differences. p < 0.05, Kruskal-Wallis and Dunn’s. **(B)** Maximum intensity projections acquired by LSFM from a sib control host (top) or a *ret*^-/-^ mutant host (bottom). Each animal was colonized with Δmot for 72 h prior to imaging. Dashed lines mark approximate intestinal boundaries.

**Figure S3.**
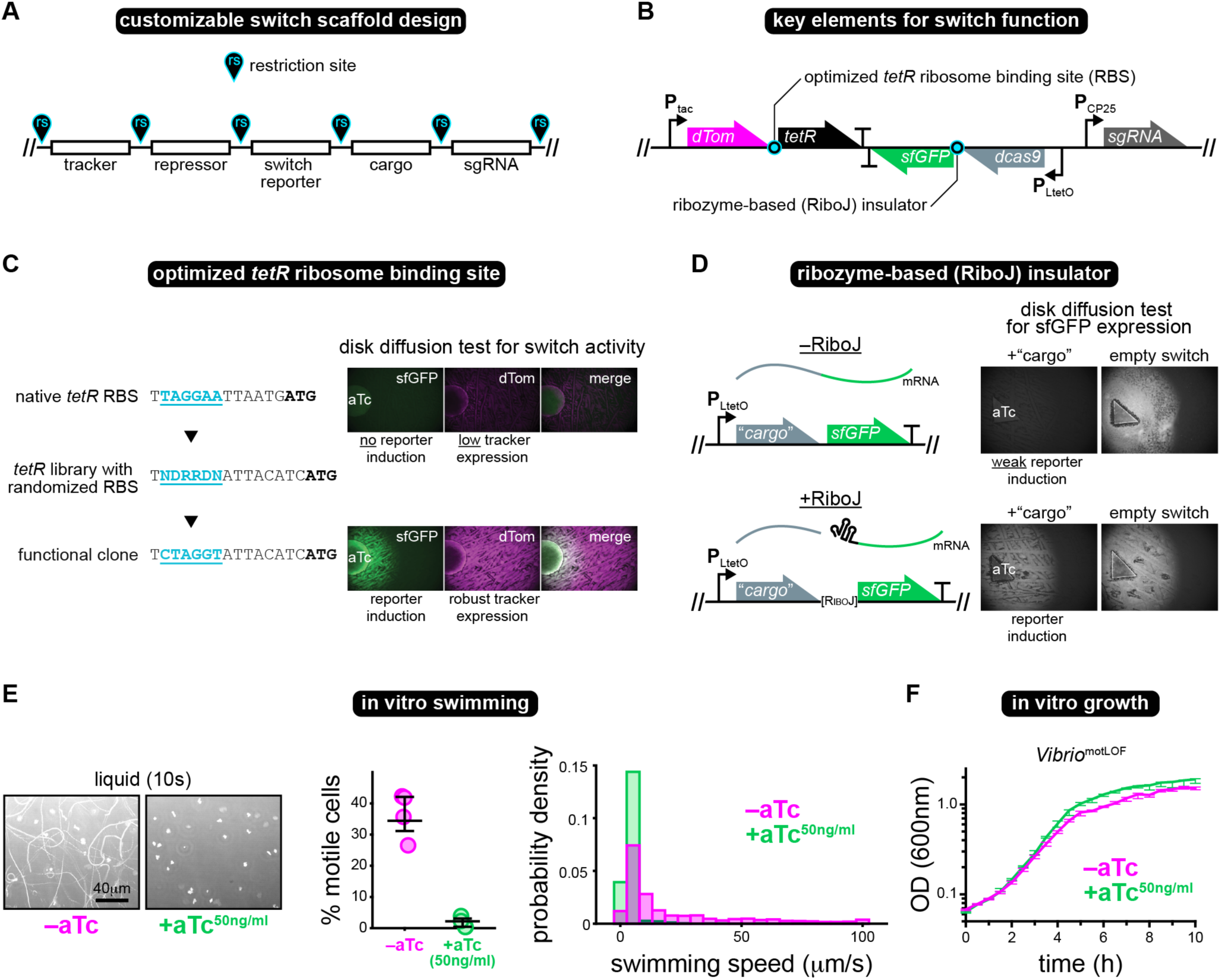
Switch design features and in vitro characterization of the motility loss-of-function switch. (associated with Figure 4) **(A)** Diagram depicts the customizable design of the switch scaffold. Unique restriction sites (rs) flanking switch elements allow each component to be optimized or replaced. The “tracker” encodes a fluorescent protein for marking all bacterial cells. The “repressor” encodes a transcription factor that allows inducible control of “cargo” gene (e.g., *dcas9*) expression. The “switch reporter” encodes a fluorescent protein that is coexpressed with the cargo gene to signal switch activation. A “sgRNA” is inserted when the switch is used for CRISPRi. **(B)** Gene diagram indicates the locations (cyan bullseyes) of two elements that were essential for switch function: an optimized *tetR* ribosome binding site (RBS) and a ribozyme-based insulator. **(C)** Left: Shown are DNA sequences for the native (top) and functionally optimized (bottom) 5’ untranslated region (UTR) of the *tetR* gene. Underlined cyan text denotes the RBS. Bolded text marks the *tetR* start codon. The middle sequence represents the library of *tetR* 5’ UTRs containing randomized RBS sequences that were screened (letters are based on IUPAC code). Right: Switch function was assessed using disk diffusion assays in which *E. coli* carrying the switch (without a cargo gene inserted) were spread at a density high enough to produce a lawn of growth on an agar plate. A disk impregnated with concentrated aTc was then used to induce switch activity, thereby making the adjacent cells express GFP if the switch was functional. Top right: Original switch prototypes failed to be induced and displayed suppressed expression of the dTomato tracker, which we surmised was due to overexpression of TetR. Bottom right: A library of switch clones containing random RBS sequences in the *tetR* 5’ UTR were screened, resulting in the recovery of a functional clone that displayed sensitive switch activation and robust tracker expression. **(D)** Top row: Early switch prototypes relied on the co-transcription of the cargo gene and sfGFP reporter. However, the insertion of large cargo genes, such as a *dcas9* or *cheA2*, hampered sfGFP expression compared to an “empty” switch without a cargo gene, which was evident in disk diffusion assays. We surmised that this was due to part-junction interference between *sfGFP* and the cargo, leading to poor translation of *sfGFP*. Bottom row: Insertion of the self-cleaving RiboJ ribozyme insulator between *sfGFP* and the cargo alleviated the apparent interference. **(E)** Left: Swimming motility of Δmot^LOF^ in liquid media plus/minus aTc (50 ng/ml). Motility in liquid media was recorded for 10 seconds on a glass slide. Images show cellular movements over the entire 10 second period, which illustrates each cell’s swimming trajectory. Motility was assessed ∼7 h post-induction. Middle: The percentage of swimming cells in Δmot^LOF^ populations in liquid media plus/minus aTc (50 ng/ml) from four separate fields of view. Right: Probability densities showing the distribution of cellular swimming speeds in liquid media for Δmot^LOF^ plus/minus aTc (50 ng/ml). Sample sizes (measured bacterial swim tracks): Δmot^LOF^-aTc = 2,677; Δmot^LOF^ +aTc = 944. **(F)** In vitro growth curves of Δmot^LOF^ in rich media (lysogeny broth) plus/minus aTc (50 ng/ml). Line traces the average optical density (OD) from three replicate wells, bars indicate range.

**Figure S4.**
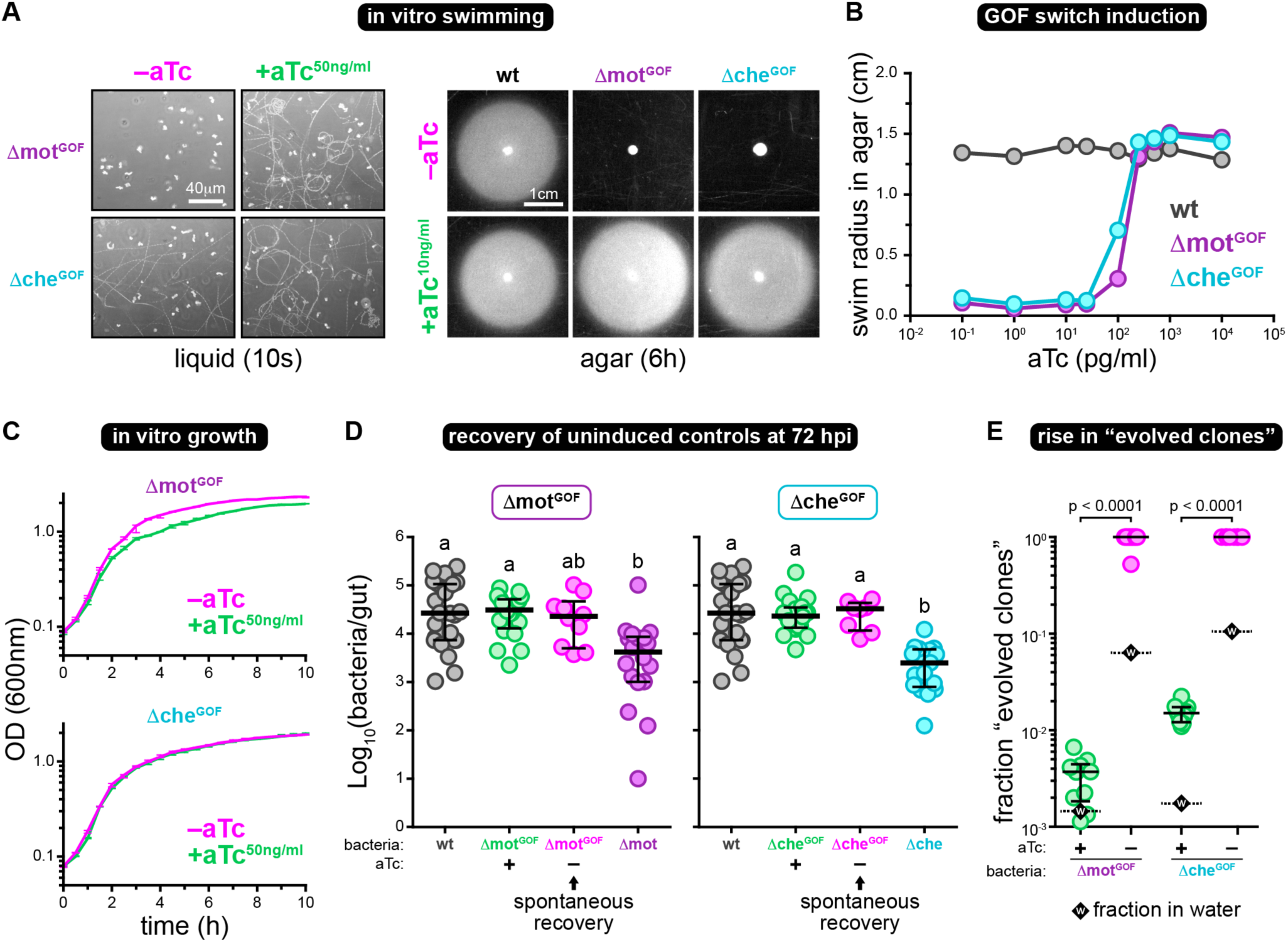
In vitro characterization of motility and chemotaxis gain-of-function switches and supporting data on the evolution of gain-of-function switches in vivo. (associated with Figure 5) **(A)** Left: Swimming motility of Δmot^GOF^ and Δche^GOF^ in liquid media plus/minus aTc (50 ng/ml). Motility in liquid media was recorded for 10 seconds on a glass slide. Images show cellular movements over the entire 10 second period, which illustrates each cell’s swimming trajectory. Motility was assessed ∼4 h post-induction. Right: Motility of wild-type *Vibrio*, Δmot^GOF^, and Δche^GOF^ in soft agar plus/minus aTc (10 ng/ml). Swim distances were captured 6 h post-inoculation of bacteria into the agar. **(B)** Swim distances of wild-type *Vibrio*, Δmot^GOF^, and Δche^GOF^ in soft agar 6 h post-induction with different concentrations of aTc. **(C)** In vitro growth curves of Δmot^GOF^ and Δche^GOF^ in rich media (lysogeny broth) plus/minus aTc (50 ng/ml). Line traces the average optical density (OD) from three replicate wells, bars indicate range. **(D)** Cultivation-based abundances of Δmot^GOF^ or Δche^GOF^ at 72 hpi either with (green) or without (magenta) aTc induction. Abundances of wild-type *Vibrio* (gray), Δmot (purple), and Δche (cyan) (from Figure 1A, 72 hpi) are shown for comparison. Abundances of each GOF strain in the presence of aTc are from Figure 5E and are also shown for comparison. Bars denote medians and interquartile ranges. Letters denote significant differences. p < 0.05, Kruskal-Wallis and Dunn’s. **(E)** Shown is the fraction of “evolved clones” (i.e., bacterial colonies recovered that displayed constitutive switch activation) from the intestines of zebrafish colonized with Δmot^GOF^ or Δche^GOF^ at 72 hpi that were either induced (green) or not induced (magenta) with aTc. Bars denote medians and interquartile ranges. In each condition, a black dashed bar and diamond labeled with a “w” indicates the fraction of evolved clones recovered from the water environment.

**Table S1.** Bacteria used and created in this study.

| Strain | Description/ Relevant Details | Source |
| --- | --- | --- |
| **Wild-type Bacteria** |  |  |
| *Vibrio* ZWU0020 | Non-toxigenic strain of *Vibrio cholerae* isolated from the zebrafish gut. IMG genome ID: 2703719078 | (Stephens et al., 2016) |
| **Recombinant Bacteria** |  |  |
| *Vibrio* Δmot | *Vibrio* ZWU0020 with unmarked, in-frame deletion of *pomAB* (non-motile) | (Wiles et al., 2018) |
| *Vibrio* Δche | *Vibrio* ZWU0020 with unmarked, in-frame deletion of *cheA2* (non-chemotactic) | This study |
| *Aeromonas* ZOR0001<br>*attTn7::dTomato* | Strain of *Aeromonas veronii* isolated from the zebrafish gut. Constitutively expresses dTomato; Gent<sup>R</sup> | (Wiles et al., 2018) |
| *Vibrio* ZWU0020<br>*attTn7::dTomato* | constitutively expresses dTomato; Gent<sup>R</sup> | (Wiles et al., 2018) |
| *Vibrio* ZWU0020 *attTn7::sfGFP* | constitutively expresses sfGFP; Gent<sup>R</sup> | (Wiles et al., 2018) |
| *Vibrio* Δmot *attTn7::dTomato* | constitutively expresses dTomato; Gent<sup>R</sup> | This study |
| *Vibrio* Δche *attTn7::dTomato* | constitutively expresses dTomato; Gent<sup>R</sup> | This study |
| *Vibrio*<sup>motLOF</sup> | *Vibrio* carrying the motility loss-of-function switch within the chromosome at the *attTn7* site; constitutively expresses dTomato; Gent<sup>R</sup> | This study |
| *Vibrio* Δmot<sup>GOF</sup> | Δmot carrying the motility gain-of-function switch within the chromosome at the *attTn7* site; constitutively expresses dTomato; Gent<sup>R</sup> | This study |
| *Vibrio* Δche<sup>GOF</sup> | Δche carrying the chemotaxis gain-of-function switch within the chromosome at the *attTn7* site; constitutively expresses dTomato; Gent<sup>R</sup> | This study |
| **Other bacteria used for molecular biology and switch optimization** |  |  |
| *E. coli* SM10 | Donor strain used for conjugation | (Simon et al., 1983) |
| DH5α | *E. coli* cloning strain | NEB |
| *E. coli* MG1655 | Used for switch prototyping and optimization | (Blattner et al., 1997) |
| *E. coli* HS | Used for switch prototyping and optimization | (Rasko et al., 2008) |
| *Enterobacter* ZOR0014 | Source of *tetR* gene | (Stephens et al., 2016) |

**Table S2.** Plasmids used and created in this study.

| Plasmid Name | Description/ Relevant Details | Source |
| --- | --- | --- |
| **Plasmids used for allelic exchange** |  |  |
| pAX1 | allelic exchange vector with GFP merodiploid tracker and temperature-sensitive replicon *ori<sub>101</sub>/repA101<sup>ts</sup>*; Amp<sup>R</sup>, Gent<sup>R</sup>, Clm<sup>R</sup>, 30°C; Addgene plasmid #117397 | (Wiles et al., 2018) |
| pAX1-ZWU0020-cheA2 | pAX1-based plasmid with *cheA2* knockout cassette; Amp<sup>R</sup>, Gent<sup>R</sup>, Clm<sup>R</sup>, 30°C; Wiles plasmid #pTW383 | This study |
| **Customizable plasmid-based genetic switch scaffolds** |  |  |
| pXS-GOF-switch | gain-of-function switch scaffold constructed in a pXS-based plasmid; Amp<sup>R</sup>; Wiles plasmid #pTW265 | This study |
| pXS-LOF-switch | loss-of-function switch scaffold constructed in a pXS-based plasmid; Amp<sup>R</sup>; Wiles plasmid #pTW308 | This study |
| **Plasmids used for Tn7-based chromosomal insertions** |  |  |
| pTn7xTS | Tn7 tagging vector with temperature-sensitive replicon *ori<sub>101</sub>/repA101<sup>ts</sup>*; Amp<sup>R</sup>, Gent<sup>R</sup>, 30°C; Addgene plasmid #117389 | (Wiles et al., 2018) |
| pTNS2 | Tn7 helper plasmid carrying transposase genes; Amp<sup>R</sup>; Addgene plasmid #64968 | (Choi et al., 2005) |
| pTn7xTS-dTomato | pTn7xTS carrying P<sub>tac</sub>-*dTomato*; Amp<sup>R</sup>, Gent<sup>R</sup>, 30°C; Addgene plasmid #117391 | (Wiles et al., 2018) |
| pTn7xTS-GOF-switch | pTn7xTS carrying the gain-of-function switch scaffold; Amp<sup>R</sup>, Gent<sup>R</sup>, 30°C; Wiles plasmid #pTW285 | This study |
| pTn7xTS-LOF-switch | pTn7xTS carrying the loss-of-function switch scaffold; Amp<sup>R</sup>, Gent<sup>R</sup>, 30°C; Wiles plasmid #pTW317 | This study |
| pTn7xTS-mot-LOF-switch | pTn7xTS-LOF-switch with *pomA* sgRNA; Amp<sup>R</sup>, Gent<sup>R</sup>, 30°C; Wiles plasmid #pTW340 | This study |
| pTn7xTS-mot-GOF-switch | pTn7xTS-GOF-switch with *pomAB*; Amp<sup>R</sup>, Gent<sup>R</sup>, 30°C; Wiles plasmid #pTW324 | This study |
| pTn7xTS-che-GOF-switch | pTn7xTS-GOF-switch with *cheA2*; Amp<sup>R</sup>, Gent<sup>R</sup>, 30°C; Wiles plasmid #pTW282 | This study |
| **Backbones and sources of genetic parts** |  |  |
| pXS-dTomato | pXS-based modular *dTomato* expression scaffold; Amp<sup>R</sup>; Addgene plasmid #117387 | (Wiles et al., 2018) |
| pdCAS9-bacteria | Source vector for *dcas9* gene; Clm<sup>R</sup>; Addgene plasmid #44249 | (Qi et al., 2013) |
| pTW168 | Source vector for *sfGFP* gene; Amp<sup>R</sup>, Gent<sup>R</sup>; Wiles plasmid #pTW168 | (Schlomann et al., 2019) |

Amp<sup>R</sup>, encodes ampicillin resistance; Gent<sup>R</sup>, encodes gentamicin resistance; Clm<sup>R</sup>, encodes chloramphenicol resistance; 30°C, permissive growth temperature

**Table S3.**
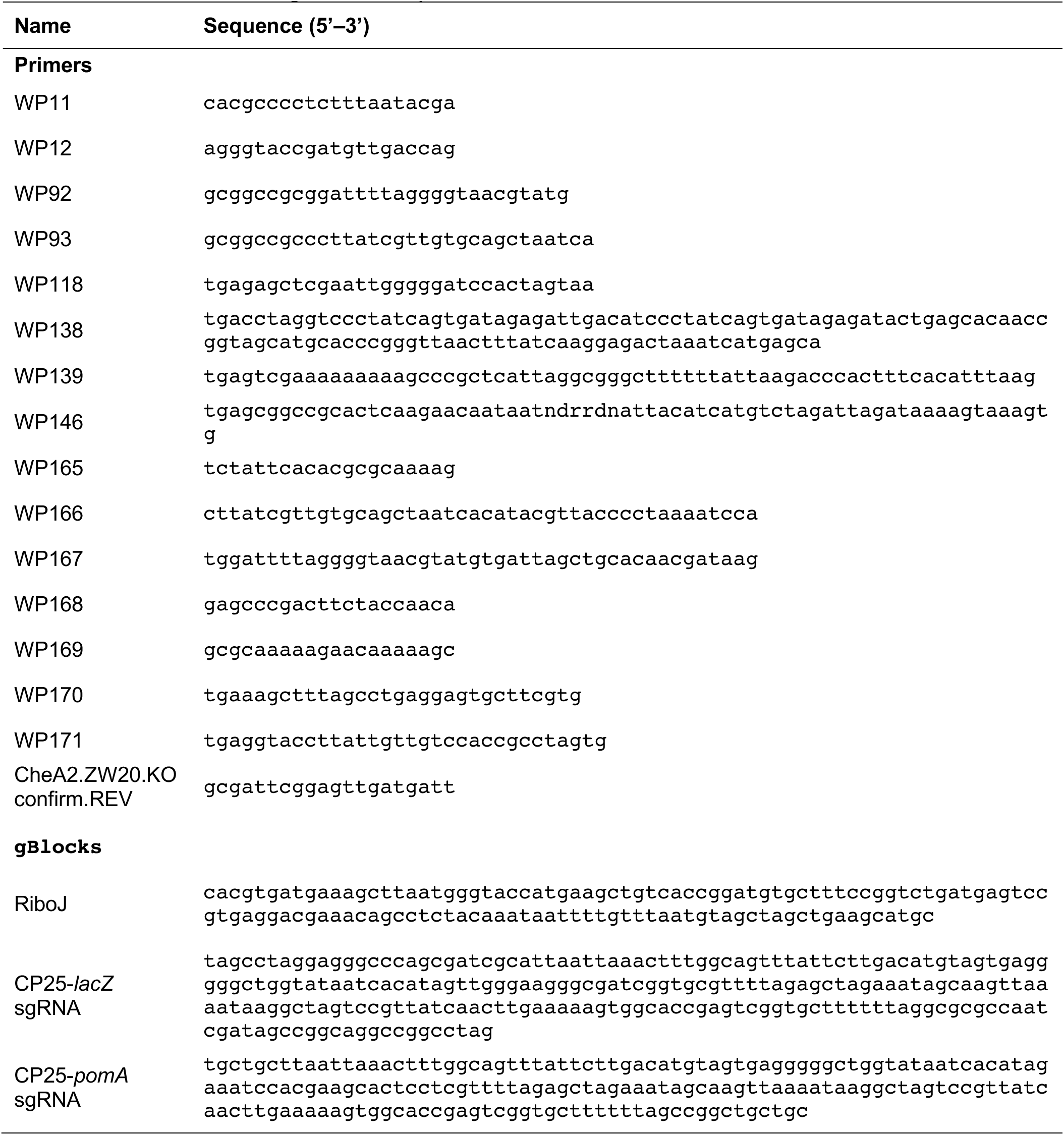
Primer and oligo DNA sequences.

